# Protein Engineering for Thermostability through Deep Evolution

**DOI:** 10.1101/2023.05.04.539497

**Authors:** Huanyu Chu, Zhenyang Tian, Lingling Hu, Hejian Zhang, Hong Chang, Jie Bai, Dingyu Liu, Jian Cheng, Huifeng Jiang

**Affiliations:** Key Laboratory of Engineering Biology for Low-carbon Manufacturing, Tianjin Institute of Industrial Biotechnology, Chinese Academy of Sciences, Tianjin 300308, P. R. China; National Center of Technology Innovation for Synthetic Biology, Tianjin, 300308, P. R. China; College of Biotechnology, Tianjin University of Science and Technology, Tianjin, 300457, China

## Abstract

Protein engineering for increased thermostability through iterative mutagenesis and high throughput screening is labor-intensive, expensive and inefficient. Here, we developed a deep evolution (DeepEvo) strategy to engineer protein thermostability through global sequence generation and selection using deep learning models. We firstly constructed a thermostability selector based on a protein language model to extract thermostability-related features in high-dimensional latent spaces of protein sequences with high temperature tolerance. Subsequently, we constructed a variant generator based on a generative adversarial network to create protein sequences containing the desirable function with more than 50% accuracy. Finally, the generator and selector were utilized to iteratively improve the performance of DeepEvo on the model protein glyceraldehyde-3-phosphate dehydrogenase (G3PDH), whereby 8 highly thermostable variants were obtained from only 30 generated sequences, demonstrating the high efficiency of DeepEvo for the engineering of protein thermostability.

## Introduction

Engineering proteins for thermostability is crucial for broadening the application of natural proteins in multiple fields such as food, feed, biocatalysis, biomedicine, and biomanufacturing^1-3^. Directed evolution is the most powerful tool for improving the thermostability of natural proteins, but it currently requires multiple rounds of random mutagenesis and high throughput screening ^2, 4-7^. However, the space of possible protein sequences is too large to search exhaustively in the laboratory or computationally, and functional proteins within the total protein sequence space are extremely scarce. As a consequence, it is very difficult to identify highly functional sequences in the vast nonfunctional sequence space^8-10^. To overcome this, many rational or semi-rational strategies^11-14^ have been developed to improve the possibility of each mutant to have the desired function, as well as many high throughput approaches^6, 7^ to increase the rate of experimental screening, but there is still a lot of room to improve the efficiency of these tools^15, 16^.

Recently, novel deep learning models have been developed to predict protein structure^17-19^, EC number^20^, enzyme turnover^21^, gene function^22, 23^, and also the thermostability of proteins^24^. Studies of protein sequence design demonstrated that deep learning models can learn the diversity of natural protein sequences and enables the generation of functional protein variants^22, 23, 25, 26^. In addition, some general protein language models, such as UniRep^27^ and ESM^28^, can encode the enormous protein sequence space into a high-dimensional representation space, in which it is more feasible to establish connections between protein properties and sequence variants^29-31^. These achievements provide us an opportunity to develop a method to engineer proteins with improved thermostability by merging two deep learning models from an iterative evolution perspective, whereby a generative model produces abundant variants from a reasonable sequence space with the desired function, after which a selective model is used to identify variants with improved thermostability.

In contrast to a typical directed evolution strategy based on highly labor-intensive iterative mutagenesis, here we proposed a deep evolution (DeepEvo) strategy to improve protein thermostability through global sequence generation and selection (Figure 1). Firstly, we leveraged a successful protein language model (ESM) to extract thermostability-related information from more than 190,000 protein sequences across a wide range of organisms, and constructed a thermostability selection model (Thermo-selector). Then, a modified generative model (Variant-generator) based on ProteinGAN was constructed to generate functional sequences. Finally, after iterative optimization of Variant-generator by the output of Thermo-selector, we evaluated the efficiency of DeepEvo for protein thermostability engineering on the model enzyme G3PDH, which is a key enzyme for glycolysis with important applications in industry and medicine^32, 33^ (Figure S1).

**Figure 1.**
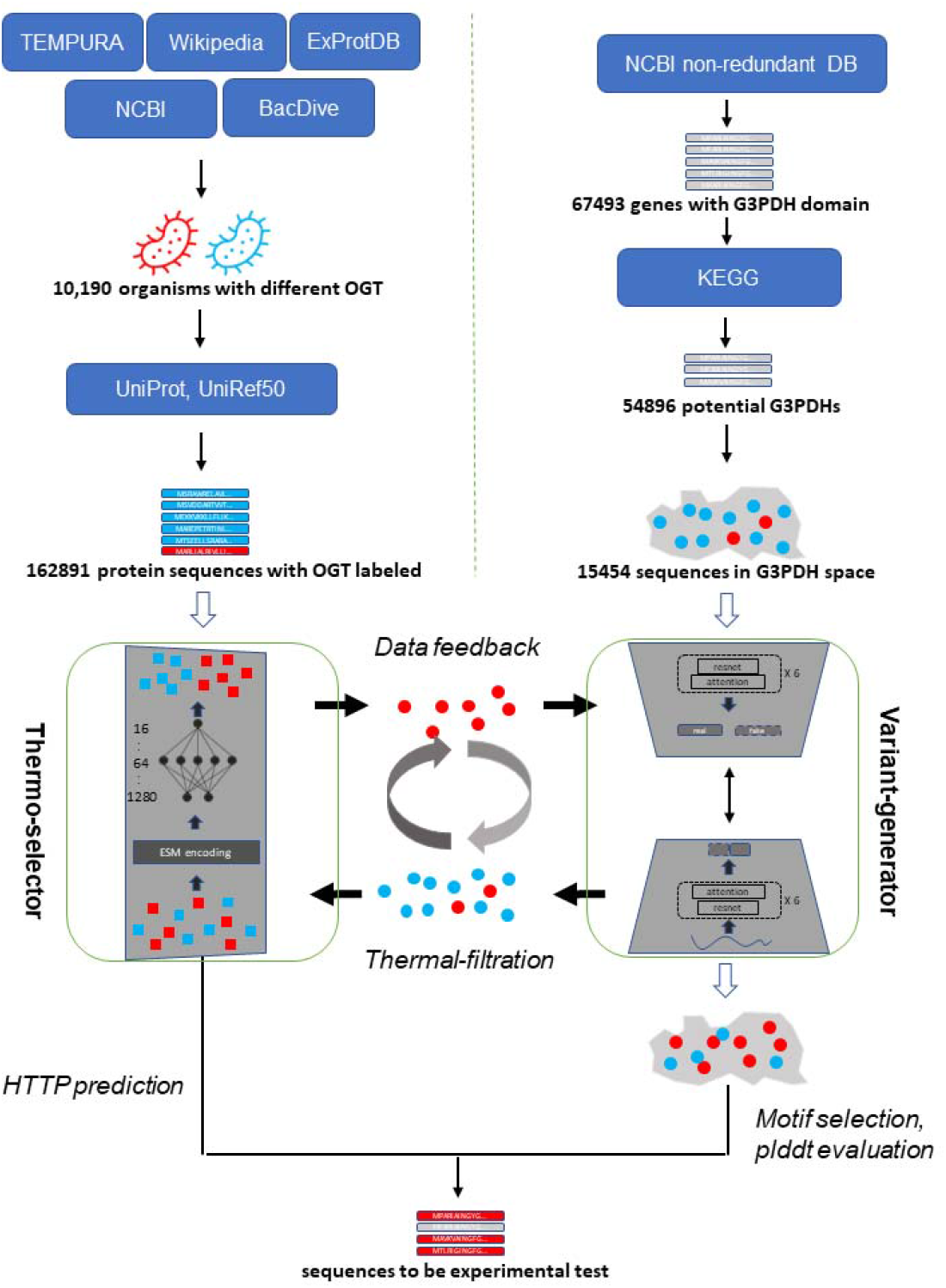
Framework and data flow of Deep Evolution. Firstly, two data collection procedures were constructed to collect natural sequences with desired function (G3PDH as example) and sequences from different optimal growth temperatures (OTG) organisms. These sequences were then used to train the Variant-generator and Thermo-selector respectively; Secondly, the Thermo-selector model was trained and used as a thermostable sequence filter to classify HTTP sequences (denoted as red squares) and LTTP sequences (denoted as blue squares), and the Variant-generator was trained to generate reasonable sequences (denoted as circles) in the confined functional protein sequence space; Thirdly, the generated sequences by Variant-generator were passed to the Thermo-selector, and the sequences predicted as HTTP (denoted as red circles) were used to refine the Variant-generator by a data feedback procedure. The generation, thermal traits filtration, feedback and regeneration constructed an iteration procedure to generate suitable sequences with high temperature tolerance directly; At last, we selected the generated sequences that predicted as HTTP and had suitable functional conservation motif and good structural predictions to perform experimental verification.

## Results

### Construction of the thermostability selection model: Thermo-selector

Similar to natural selection, the DeepEvo approach utilized a thermostability selection model to identify protein sequences with potential for enhanced thermostability. A thermostability selection model was constructed to predict whether a protein is a high temperature tolerant protein (HTTP) or low temperature tolerant protein (LTTP). Considering that the total proteins of organisms that survive in high-temperature environments should be HTTP, the optimal growth temperatures (OGT) of the organisms were used as a label to measure the thermostability of the natural protein. To obtain enough labeled data for supervised learning, the information of 10,190 organisms with a wide range of OGT was collected from TEMPURA^34^, ExProtDB^35^, NCBI, and BacDive^36^. Then, more than 20 million corresponding genes were retrieved from UniProt and UniRef gene sets. The corresponding protein sequences from organisms with OGT > 50 □ or OGT < 30 □ were defined as HTTPs or LTTPs, respectively (Figure1, Methods). To reduce the impact of sequence similarity on the thermostability-related traits, only sequences with pairwise identity less than 50% were retained. In view of the length of most enzymes used in practical applications, proteins with a length > 300 and < 800 amino acids were retained. After filtering based on these criteria, a total of 30,968 HTTPs and 162,890 LTTPs were collected to build the selection model (Figure1, Table 1).

**Table 1.**
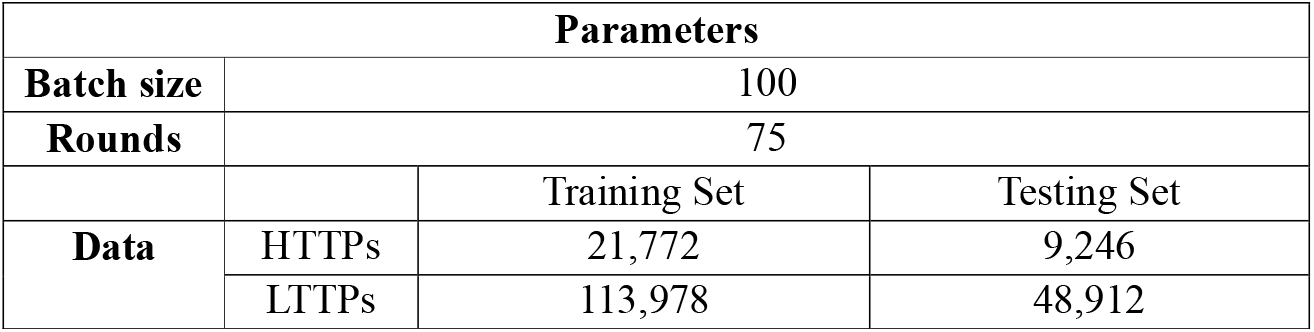
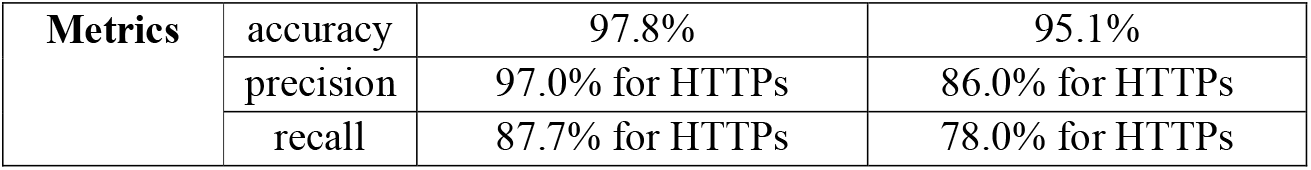
Summary of the thermo-selector model.

Inspired by natural language processing techniques, the ESM-1b pre-training model was used to encode the training data as a 1280-dimensional vector. Using the ESM embedding vectors as input, a three-layer fully connected neural network was built (Table 1). Using 70% of the collected data as the training set, the model was optimized by cross-entropy loss. After 75 rounds of training, the loss function of the model became stable (Figure S2). After the training procedure, the overall accuracy of the model on the testing set comprising the remaining 30% of the data was 95.1% (Figure S3), indicating that most of proteins in the testing set were correctly classified into HTTPs or LTTPs. Considering the imbalance of our training set, with 84% of total sequences belonging to LTTPs, we calculated the precision and recall to further evaluate the performance of our model on the tested HTTPs (Supplementary Methods). These measurements showed that 86.0% (precision) of all labeled HTTPs were predicted as HTTPs by the model, and 78.0% (recall) of all sequences predicted as HTTPs by the model were actually the labeled HTTPs in the test set (Table 1). These results suggest that our model can be used as a viable filter for identifying HTTPs. This thermostability selection model was named Thermo-selector.

### Construction of a variant generation model for G3PDH: Variant-generator

To more efficiently generate functional sequences in a confined sequence space by DeepEvo approach, a G3PDH Variant-generator was built by revising ProteinGAN with the multi-headed attention mechanism (Figure 1 and S4). This model structure includes a sequence generator and a discriminator. The generator attempts to generate functional sequences and the discriminator attempts to distinguish the generated sequences from the natural sequences. By searching with G3PDH functional domain and filtering with sequences length and identity, 15,454 natural G3PDH sequences were extracted from the NCBI, KEGG, and Pfam databases to train the model (Methods). At each training step, starting from a random vector, the generator produced 64 sequences, which were mixed with the same number of natural G3PDH sequence. The discriminator then compared the generated sequences with the natural sequences, which were used to adjust the parameters of both the generator and the discriminator. After 200,000 training rounds, the sequences produced by the generator could not be distinguished from the natural G3PDH sequences by the discriminator (Figure S5 and S6).

To evaluate the quality of these generated sequences, we conducted t-distributed stochastic neighbor embedding (t-SNE) dimensionality reduction on the natural and generated G3PDH sequences (Figure 2A, left pane). The generated sequences covered a similar distribution to that of the natural sequences, and were grouped into smaller clusters and interpolated within the natural sequence clusters, indicating that the Variant-generator model expanded the sequence space of natural G3PDHs. To verify the evolutionary properties reflected in the statistics of amino acid variation, we computed Shannon entropies for each position in multiple sequence alignments of the generated and natural G3PDH sequences. The positional variability of the generated sequences was highly similar to that of the natural sequences (Figure S7). We also evaluated the highly conserved regions related to the function of G3PDH and found that the generated sequences captured these key positions faithfully (Figure 2B).

**Figure 2.**
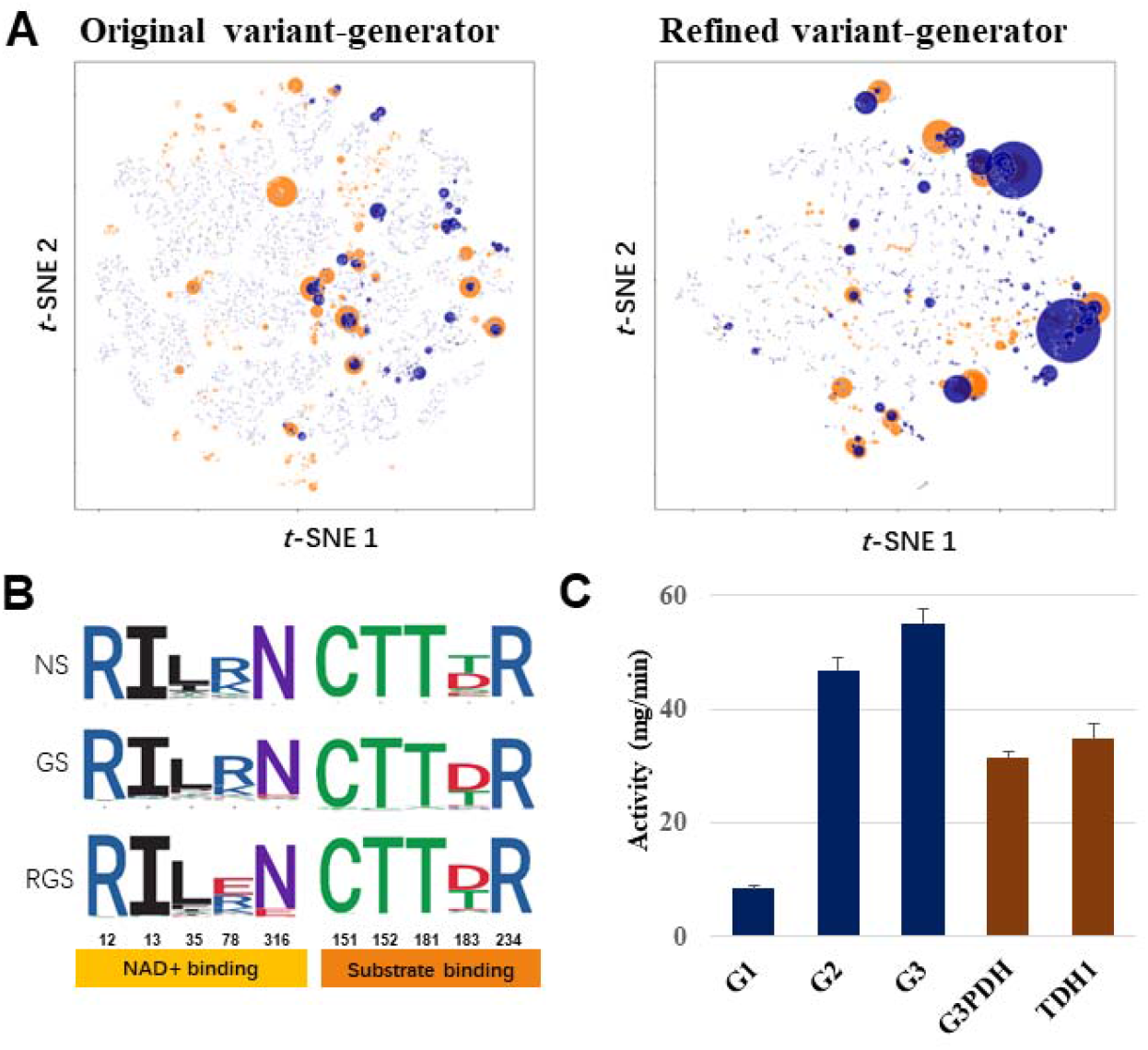
Evaluation of the sequences generated by Variant-generator. **(A)** t-SNE maps of generated sequences by original Variant-generator and refined Variant-generator. Sequences were classed in the 2-dimentional t-SNE space. The natural sequences clusters are shown as orange circles and the generated sequences clusters are shown as blue circles. The area of the circles indicates the size of the clusters. **(B)** Sequence logos of binding pockets of natural sequences (NS), original generated sequences (GS) and refined generated sequences (RGS). The conserved positions are grouped in NAD+ and substrate binding. **(C)** The activity of G3PDHs generated by the original Variant-generator at 30□. G1, G2 and G3 represent the three generated sequences, a commercial G3PDH from *rabbit* and the G3PDH from *yeast* (TDH1) were used as control.

To further evaluate the function of the generated sequences, we sorted them based on the score of the discriminator and filtered them based on the key functional conserved sequence motifs (Methods). Then, 10 sequences with different similarities to the natural sequences were selected as input for alphafold2 to build protein structure, and 6 sequences (G1-G6) with high plddts (>90%) were selected for further experimental validation. Among the 6 proteins, three (i.e., G1, G2, G3) not only folded correctly in *E. coli* expression systems (Figure S8A), but also displayed normal G3PDH activity in *in vitro* (Figure 2C). G2 and G3 even showed higher activities than the natural G3PDH from yeast and a commercial G3PDH from rabbits. These experiments proved that the Variant-generator could efficiently generate functional variants from the confined enzyme sequence space.

### Development of the deep evolution process

Based on the good performance of Thermo-selector and Variant-generator, we further implemented the DeepEvo strategy by iterating the two models to enhance sampling in the G3PDH functional sequence space for variants with enhanced thermal stability (Figures 1 and S4). First, 18,238 sequences were selected from the 100,000 generated sequences of the initial Variant-generator based on the discriminator score and the functional conserved residues of G3PDH. Then, the selected sequences were input into the Thermo-selector, where only 1,354 (7.4%) variants were classified as HTTPs. Finally, 1,354 HTTPs were mixed with all natural HTTPs and added back to the training set of the Variant-generator to refine the model. When the Variant-generator stabilized again, we obtained a refined Variant-generator, which displayed a better HTTP generation performance, as the proportion of HTTPs among the generated sequences increased to 14.9%. Additionally, using the discriminator score as metric, we observed that the sequences generated by the refined Variant-generator were largely consistent with those of the initial variant-generator (Figure S6). Interestingly, the t-SNE analysis of the new generated sequences yielded some bigger clusters, which suggested that the generated sequences were enriched in sequence spaces, similar to gene family evolution in nature, indicating that the iterative process of DeepEvo might recapitulate certain unsought mechanisms of the natural evolution process (Figure 2A, right pane).

To further evaluate the thermal stability of the sequences generated by the refined Variant-generator, 30 sequences (G7-G36) were selected from 2760 newly generated HTTPs for experimental validation based on the discriminator score, conserved residues, similarity to the nearest natural sequences and plddts (Methods). These sequences exhibited an average 61% sequence identity among themself (Figure S9), and a range of identities (∼70 to ∼90%) to their nearest natural sequences in the training set (Supplementary Table 2). The 30 selected sequences were synthesized and then expressed in *E. coli* for protein purification. Among the 30 proteins 23 (77%) were soluble and could be purified (Figures S8B-D), 17 of which (57%) showed normal G3PDH catalytic activity at 30 □ in the subsequent G3PDH activity assay (Figure 3A, Supplementary Table 2). We found that 11 out of the 17 proteins showed detectable activity at 65 □, with 8 of them (i.e., G7, G8, G10, G11, G12, G13, G14, G15) exhibiting relatively high thermostability (Figure 3A, Methods), even retaining activity at 70 and 75 □ (Figure S10). Notably, the nearest natural homologs of 7 among the 8 proteins showed low or even no detectable activity at 65 □ (Figure 3B), even one of them(N12) was from a high temperature organism, indicating that DeepEvo indeed can effectively engineer natural LTTPs into HTTPs.

**Figure 3.**
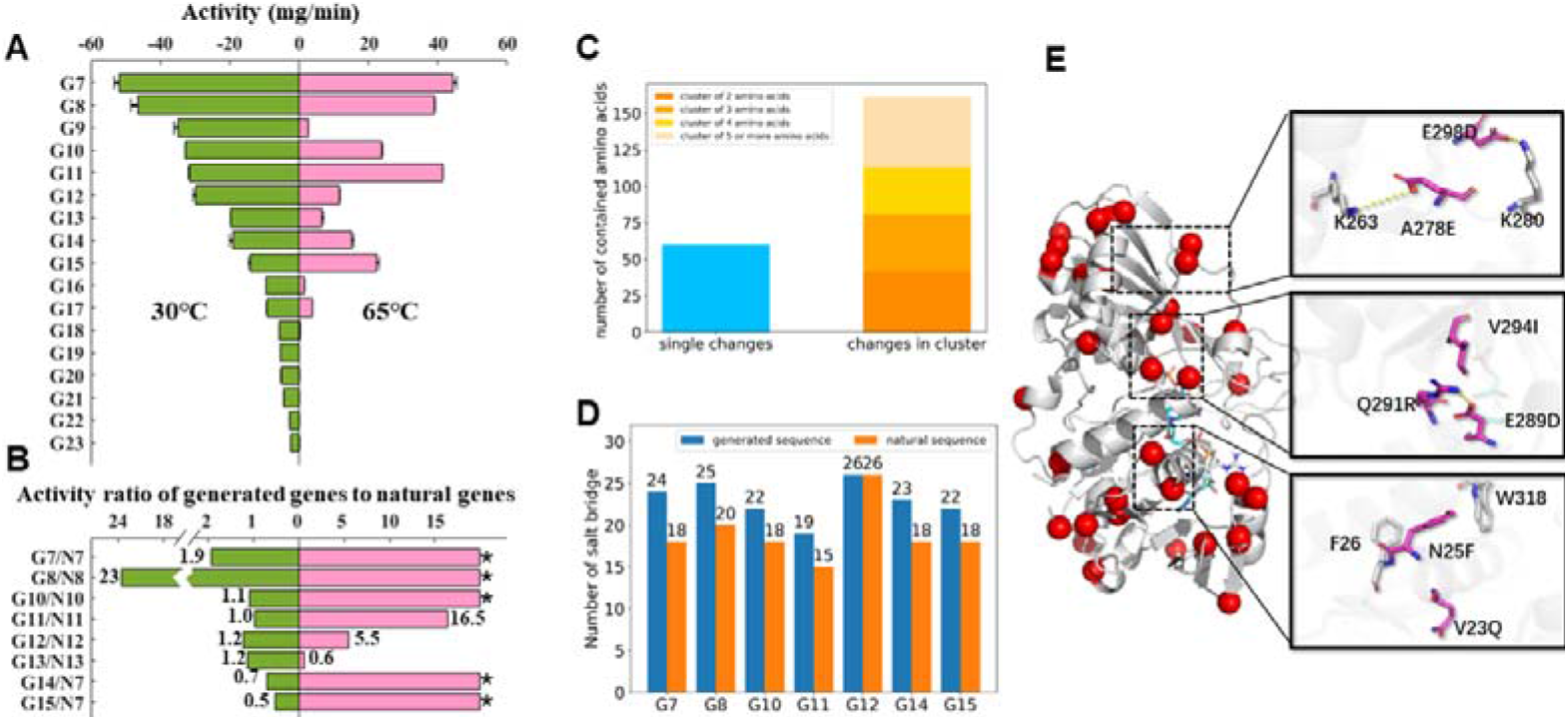
Evaluation of the thermostable G3PDH generated through DeepEvo. **(A)** Catalytic activities of 17 generated G3PDHs at 30□ and 65□ respectively. **(B)** Activity ratio of eight verified HTTPs to their individual nearest natural sequence. ‘*’ refers to no activity was detected in natural sequence, and the denominator is 0 in ratio value calculation. **(C)** Histogram of single changes and clustered changes of mutations in 7 high temperature activity significantly improved variants. Clusters contains different number of amino acids are shown in different color. **(D)** Salt bridge numbers of verified HTTPs compared with the number of there corresponding natural proteins. The numbers are calculated by ESBRI^37^. **(E)** Structure model of G8 and the single point mutations (red dot) compared to N8. Substrate and coenzyme are shown as blue sticks. Three clusters of mutations are shown in right black boxes.

### Deciphering the design art of deep evolution

In order to comprehensively understand the design art of DeepEvo, we compared 7 generated HTTPs with their nearest natural LTTPs, finding that the natural LTTPs require the mutation of approximately 20-50 residues to become our generated HTTPs (Supplementary Table 2). We observed many alanine to serine mutations, which may increase the coordination of local hydrogen bonding networks. In addition, we found that a high proportion of mutations introduced charged residues, resulting in a significant increase in the number of salt bridges^37^ in most of the verified HTTPs (Figure 3D), which could strengthen the local residual interactions and may be a major reason for the stability of the generated HTTPs^38^. The remaining mutations mainly introduced the same type of amino acid (Figure S12), and generally did not have particularly strong effects on the local side-chain arrangements. Structural analysis showed that the mutated amino acid resides were mainly distributed on the protein surface, with only a few occurring near the catalytic pocket (Figure S11). Interestingly, we found that 2/3 of the point mutations formed spatial clusters or mutation networks, in which mutants with at least one backbone alpha carbon atom (CA) are within 8Å of the CA of other mutants. Conversely, only 1/3 of the point mutations were single changes (Figure 3C and Supplementary Table 3). These results suggest that DeepEvo can enhance local structural interactions and compensate for the deleterious effects of single point mutations through the interaction of multiple mutation sites.

In order to scrutinize the interaction of spatial clusters, we compared the protein G8 with its nearest natural sequence N8, since it showed a great increase of both enzyme activity and thermal stability (Figures 3A and B). We observed a total of 34 residue changes and 25% more electrostatic interaction pairs in G8 than in N8 (Figure 3D), which may contribute the overall improved stability of G8 at high temperature according to MD simulations (Figure S13). Among these mutations, approximately 65% (22/34) were located in 7 spatial clusters (Figure 3E). For example, the A278E mutation in cluster 1 added a new salt bridge, which could change the local position of the adjacent K280. In order to keep the original salt bridge with K280, DeepEvo made the additional mutation E298D (Figure 3E top). Similar to cluster 1, the mutation Q291R in cluster 2 would add a pair of salt bridges, but two extra mutations (E289D and V294I) occurred nearby, compensating for the effect of changed residue volume (Figure 3E middle). Different from clusters 1 and 2, an enhancement in local hydrophobic stacking was observed in cluster 3, in which a π-π interaction was added to strengthen the interaction between the helix bundle through N25F and a nearby residues, while a V23Q mutation might compensate for the increased solvent exposure in the opposite direction (Figure 3E, bottom). These results indicate that the algorithm did not simply increase local interactions, but also changed the surrounding residues in clusters to achieve a more reasonable local structure, which is often a challenge for conventional enzyme engineering ^39, 40^. Thus, the DeepEvo strategy, using the Variant-generator to consider the context of residues, may enable much deeper sampling in the confined functional sequence space. This new design art, which relies on the synergistic action of multiple mutant sites, may be useful in overcoming local optima.

## Discussion

Here we present the novel protein engineering strategy DeepEvo based on deep learning models (Figure 4). Similar to directed evolution, the iteration process is key to the efficiency of the DeepEvo strategy^5, 41, 42^. When we tested the performance of Thermo-selector on generated sequences without an iteration procedure, only 4 out of 30 generated G3PDHs exhibited activity at 65 □ (Figure S14). The iteration procedure, which uses the generated HTTPs screened by the Thermo-selector to refine Variant-generator, accumulates thermostable traits in a process similar to natural evolution^43^. Our results indicated that feedback and regeneration improved the proportion of experimentally tested HTTPs among the generated sequences and compensated for the data limitation. Overall, 11 out of 30 G3PDHs generated by the algorithm (Supplementary Table 1, G7-G36) showed activity at 65 □. In addition to G3PDH, we also successfully obtained highly thermotolerant variants of malate dehydrogenase (MDH) (Figure S15), which has been used for the evaluation of multiple protein language models^22, 23^. With the development of next-generation methods^44, 45^, more rounds of iteration and more valuable thermotolerance-related data can be applied to optimize the whole DeepEvo process.

**Figure 4.**
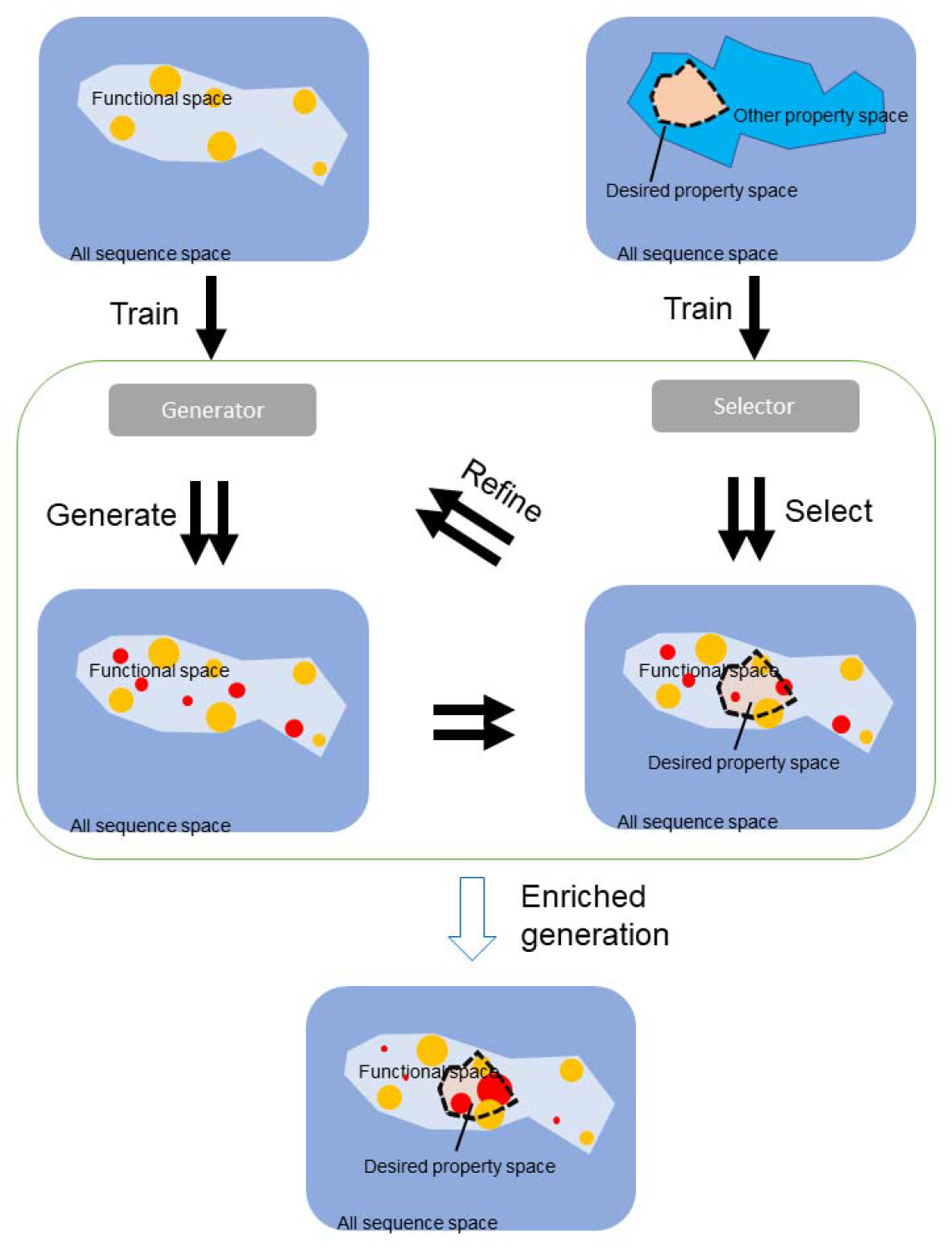
DeepEvo in the perspective of sequence space. In the vast full sequence space of proteins, the functional sequence space occupies only a small fraction. Natural sequences cluster like islands in the functional space (orange circles). The DeepEvo strategy can sample much deeper into the functional space using a trainable generator that fills in the gaps between natural sequence islands (rad circles). For particular desired properties, we can train special selectors to filter the generated sequence into the desired property space and refine the generator. After iterations, the sequences sampled by DeepEvo can be enriched in the desired property space, improving the efficiency of obtaining proteins with desired properties.

Billions of years of natural evolution have produced an immeasurable wealth of functional proteins, which nevertheless occupy only a tiny fraction of the practically endless potential protein sequence space^8, 46^. Directed evolution, similar a boat cruising around an island in a vast unexplored sea, only locally searches for beneficial mutants around natural proteins by iterative mutagenesis and high throughput screening^47^. However, the complete landscape of functional proteins contains “cliffs” and “holes” where small changes in sequence might result in complete loss of function^48^. By enabling us to obtain a better understanding of the whole landscape of protein diversity, DeepEvo is accessible to acquire previously unexplored sections of the potential sequence space. This strategy reduces the likelihood of generating non-functional sequences, thereby improving the screening efficiency (Figure 4). By concentrating on the relatively small functional sequence space and employing a thermostable selector for feature enrichment, our method significantly boosts the screening efficiency. Most of the HTTPs we generated had more than 20 mutations when compared to their closest natural sequences, which would result in theoretically trillion-level combinatorial libraries that make experimental or computational screening challenging^49^. However, our method generates variants in the reduced functional space constrained by a specific desirable property that circumvents the issue of effectively combining single-point mutations, making it highly applicable in the field of protein engineering.

In summary, DeepEvo employs an iteration process consisting of generation and selection to effectively produce protein sequences that possess strong foldability and high-temperature tolerance. In the future, it is possible to apply DeepEvo for engineering other protein properties such as acid-base tolerance and antigen affinity^16^, allowing for the generation of new proteins with diverse desired properties. Furthermore, we aim to explore the integration of generative frameworks from the fields of natural language processing and image processing to enhance the sequence generation results. This will further expand the potential of protein engineering through our DeepEvo approach.

## Methods

### Collection of thermophile organisms

We collected thermophile organisms (mainly microorganism) from five sources. The first source is TEMPURA which is a database of growth temperatures of usual and rare prokaryotes (http://togodb.org/db/tempura). In the database, we obtained about 8000 organisms and their optimal growth temperatures (OGT); The second source is ExProtDB, which is a database collecting extremophilic proteins and their host organisms, about 300 thermophiles were collected from this database; The third source was Wikipedia web search. We fetched the names of all genome sequenced microorganisms from NCBI, and then searched the name in the web. If the web contains some key words like ‘extremophile’, ‘thermophilc’, ‘thermophile’, ‘thermophilic’, ‘high temperatures’, ‘thermoacidophilic’ or ‘polyextremophile’, we check the microorganism in the web whether is a thermophile organism, and about 500 thermophiles were collected by this way; The last source is BacDive which represents a collection of organism-linked information covering the multifarious aspects of bacterial and archaeal biodiversity. We collected about 5,000 microorganisms in the database which includes the information of growth temperatures. In totally, we collected 10,190 organisms, some of which without the information of OGT were individually searched in website. Among them, we define 805 organisms (with OGT >=50 □), 5122 organisms (with OGT >=30 □, <50 □) and 4262 organisms (with OGT < 30 □) as high temperature organisms (HTO), middle temperature organisms (MTO) and low temperature organisms (LTO), respectively.

### Collection of thermophile genes

For the collected about 10,000 microorganisms with the information of growth temperatures, we respectively fetched the corresponding genes from three downloaded gene sets (i.e., UniProt reference proteomes, UniRef90 and UniRef50). The fetched genes were further divided into HTO, MTO, and LTO genes. In UniProt reference proteomes, we totally obtained 25,724,264 genes which include 1,393,345 HTO, 12,317,734 MTO and 12,013,185 LTO genes, respectively. In UniRef90, we totally obtained 15,901,817 genes which include 973,655 HTO, 7,941,331 MTO and 6,986,831 LTO genes, respectively. In UniRef50, we totally obtained 2,199,998 genes which include 165,625 HTO, 1,120,580 MTO and 913,793 LTO genes, respectively. These genes were considered as training set for the construction of high temperature discrimination model.

### Collection of glyceraldehyde 3-phosphate dehydrogenase (G3PDH) genes

To construct gene generative model, we screened and analyzed all potential G3PDHs in the NCBI database. First, we predicted all potential G3PDHs by search against the non-redundant database with the Pfam domain ID PF02800 (hmmscan --cpu 10 -- domtblout output.txt -E 1e-4 PF02800.hmm NR.fasta), 67,493 genes with the domain were obtained. Second, we retrieved all G3PDHs from KEGG database (https://www.genome.jp/dbget-bin/get_linkdb?-t+genes+pf:PF02800). Third, we made a local blastp search using G3PDHs from NCBI as the query sequences, and G3PDHs from KEGG as the BLAST database. After blastp search, 54,896 potential G3PDHs were screened with three standards: the best hit is a glyceraldehyde 3-phosphate dehydrogenase (EC:1.2.1.12, KO: K00134), the identity is more than 40, and the align length is more than 200. After filtering too long and too short genes, 40,000 potential G3PDHs were selected to construct generative model.

### Building and training the Thermo-selector

To identify sequences of high temperature resistant proteins, a two-part dataset was compiled, consisting of 30,968 high temperature sequences (OMT >= 50) and 162,890 low temperature sequences (OMT < 30). These sequences are range from 300 to 800 amino acids length and the sequence identity between each other are no more than 50%. 70% of the sequences were chosen as training set and others remained as testing set. The training data was first preprocessed by encoding each sequence into a 1280-dimensional vector using the pre-trained ESM-1b model. These encoded vectors were then used to train a three-layer multilayer perceptron with dimensions 1280:64:16 and a binary cross-entropy loss function. An Adam optimizer was used to train the model with a learning rate 1 × 10^−3^. 75 epochs of training were performed to make the loss stable. The model was evaluated using standard metrics such as precision, recall, and F1 score (Supplementary Methods). The pytorch framework was used for building this model.

### Building and training the Variant-generator

To build the Variant-generator, we filtered the collected G3PDH sequences with the length > 300 and < 800 amino acids. A total of 15,454 sequences were used for training and testing. We randomly split these sequences in the ratio of 9:1 as the training set and test set respectively. The GAN architecture to generate G3PDH sequences was based on the ProteinGAN model. The discriminator and generator networks were built by ResNet blocks which contained three convolution layers with rectified linear unit activations and a transformer block with muti-head attention mechanism. A random vector of 128 values was used as the input to the generator, and the output matrix dimensions were 512 × 21, which was correspond to the one-hot encoded sequence of length 512 with a 21 words vocabulary (20 amino acids and a sign for gaps at the beginning or ending of the sequence). The matrix with the same dimensions as the output of the generator is used as input to the discriminator. In the training process, the generator generated 64 sequences as a batch, and these generated sequences were mixed with 64 natural G3PDH sequences sampled in the training set based on the sampling weights described above, and then they were passed to the discriminator for discrimination. A non-saturating loss with R1 regularization was used as loss function in this model, and we selected the Adam algorithm for optimizing the networks. The learning rate was gradually decreased from 1 × 10^−3^ to 5 × 10^−5^. The model was trained for 200,000 steps, which took about 12 hours on a Nvidia GTX2080Ti system.

### Analysis of generated sequences

A distance matrix of cluster representatives was used as the t-SNE input. To obtain cluster representatives, the numbers of sequences in both datasets were first equalized by taking 10000 sequences from natural and generated datasets. These sequences were independently clustered using MMseqs2 with 80% minimal sequence identity. Representative sequences of these clusters were chosen based on the MMseqs2 output. From the representative sequences, a distance matrix was generated using Clustal Omega. The distance matrix was used with the scikit-learn t-SNE module with default settings, except that the embedding generation perplexity was set to 7. Coordinates given by t-SNE were used for plotting and the size of a given dot was visualized based on the cluster size it represents.

### Select generated sequences for experimental test

To filter out representative sequences for experimental testing, we first used the discriminator of the GAN-based Variant-generator. After ranking the generated sequences by this score, the top 20% that were strongly discriminated as natural-like sequences were kept. We then used a crystallized G3PDH (PDBID: 3KV3) as a template, extracted the NAD and G3P binding positions (residue numbers 12, 13, 35, 78, 316 for NAD binding, 151, 152, 181, 183, 234 for G3P binding), and constructed a functional motif. We then aligned the generated sequences to the template, and if there was no gap in the functional motif region, the sequences were retained. We then calculated the identities of the generated sequences and the natural sequences with blastp. Tens of sequences were selected with different levels of variation (60%-90% for the original Variant-generator and 80%-90% for the refined Variant-generator). The selected sequences were then structurally modeled with Alphafold2, keeping those with plddt >90%.

### Expression and purification of Proteins

Protein coding DNA sequences mentioned in this study were all synthesized, cloned into pET28a expression vector between NdeI and XhoI then sequence-verified by Zhong He Gene Co.Ltd (Tianjin). The constructs were transformed into BL21 (DE3) or Arctic Express (DE3) *E. coli*. Cells were seeded in 2YT medium (kanamycin, 50 μg/mL) at a ratio of 1:160 and grown at 37 °C, 220rpm. After OD_600_ of cells were reaching 0.4∼0.6, IPTG was added to a final concentration of 0.5□mM IPTG to induce expression. Strain cells were cultured at 16 °C, 220 rpm overnight then harvested by centrifugation. Cells were resuspended in lysis buffer (50mM Tris-HCL, pH6.8) and lysed by using a high-pressure homogenizer at 1200∼1500 bar, for 2-3 times. Cell debris was discarded by centrifugation at 10,000□× g for 40□min. The Ni-NTA agarose column was balanced with ddH_2_O and lysis buffer for 2 column volume. The supernatant was applied to the column then proteins were eluted using a gradient of elution buffer (50mM Tris-HCL with 10mM, 50mM 200mM imidazole). The fractions were then collected and analyzed by SDS-PAGE. Purified proteins were concentrated by centrifugation (4,000g, 30□min) in 10□kDa ultrafiltration tubes (Centriplus YM series, Millipore) and flash frozen in liquid nitrogen then stored at - 80 °C.

### G3PDH activity and thermal stability assay

The assay for G3PDH activity was carried out according to the method originally described by Ferdinand^50^ with minor modifications. Briefly, the activity can be monitored by measuring the formation of NADH. Triplicate samples of purified proteins were mixed with 10mM NAD in 993uL Reaction (40mM triethanolamine, 50 mM Na_2_HPO_4_, 5 mM EDTA, 0.1 mM DTT, pH8.6) separately. 7 μl DL-G3PDH solution(sigma) were added into systems, then determining the A_340_ immediately. The reaction systems were incubated at 30 °C for 10 min and determining the A_340_ again.

The activity of G3PDHs were calculated with the formula Units/mg/min = △A_340_ x VT(Volume of tube)/(6.22 x Concentration(mg) x Time(s)). For the thermal stability assay, 100 μl reaction systems were developed in 96 well plate. The plates were incubated at designed temperature in a thermostable microplate reader with persistent reading of A_340_ for 30 min.

## Author contributions

All authors listed have made a substantial, direct and intellectual contribution to the work, and approved it for publication.

## Funding

This Research Topic was supported by the National Key R&D Program of China (2022YFC2106000), Tianjin Synthetic Biotechnology Innovation Capacity Improvement Project (TSBICIP-KJGG-009-02, TSBICIP-C□RC-015, TSBICIP-CXRC-003), the CAS Project for Young Scientists in Basic Research (YSBR-072-4).

## Acknowledgements

We thank all the contributing authors and reviewers for their support in this Research Topic.

## Conflict of interest

The authors declare that the research was conducted in the absence of any commercial or financial relationships that could be construed as a potential conflict of interest.

## Notes

### Competing Interest Statement

The authors have declared no competing interest.

## References

1. Bommarius, A.S. & Paye, M.F. Stabilizing biocatalysts. Chem Soc Rev 42, 6534–6565 (2013).

2. Bell, E.L. et al. Directed evolution of an efficient and thermostable PET depolymerase. Nat Catal 5, 673–681 (2022).

3. Liu, Q., Xun, G. & Feng, Y. The state-of-the-art strategies of protein engineering for enzyme stabilization. Biotechnology advances 37, 530–537 (2019).

4. Sun, Z., Liu, Q., Qu, G., Feng, Y. & Reetz, M.T. Utility of B-Factors in Protein Science: Interpreting Rigidity, Flexibility, and Internal Motion and Engineering Thermostability. Chem Rev 119, 1626–1665 (2019).

5. Packer, M.S. & Liu, D.R. Methods for the directed evolution of proteins. Nature Reviews Genetics 16, 379–394 (2015).

6. Markel, U. et al. Advances in ultrahigh-throughput screening for directed enzyme evolution. Chem Soc Rev 49, 233–262 (2020).

7. Zeng, W.Z., Guo, L.K., Xu, S., Chen, J. & Zhou, J.W. High-Throughput Screening Technology in Industrial Biotechnology. Trends in Biotechnology 38, 888–906 (2020).

8. Hie, B.L. et al. Efficient evolution of human antibodies from general protein language models. Nat Biotechnol (2023).

9. Yang, K.K., Wu, Z. & Arnold, F.H. Machine-learning-guided directed evolution for protein engineering. Nat Methods 16, 687–694 (2019).

10. Qu, G., Li, A., Acevedo-Rocha, C.G., Sun, Z. & Reetz, M.T. The Crucial Role of Methodology Development in Directed Evolution of Selective Enzymes. Angew Chem Int Ed Engl 59, 13204–13231 (2020).

11. Sun, Z., Wikmark, Y., Backvall, J.E. & Reetz, M.T. New Concepts for Increasing the Efficiency in Directed Evolution of Stereoselective Enzymes. Chemistry 22, 5046–5054 (2016).

12. Leman, J.K. et al. Macromolecular modeling and design in Rosetta: recent methods and frameworks. Nature methods 17, 665–680 (2020).

13. Delgado, J., Radusky, L.G., Cianferoni, D. & Serrano, L. FoldX 5.0: working with RNA, small molecules and a new graphical interface. Bioinformatics 35, 4168–4169 (2019).

14. Gumulya, Y. et al. Engineering highly functional thermostable proteins using ancestral sequence reconstruction. Nat Catal 1, 878–888 (2018).

15. Huang, P. et al. Evaluating protein engineering thermostability prediction tools using an independently generated dataset. ACS omega 5, 6487–6493 (2020).

16. Chautard, H. et al. An activity-independent selection system of thermostable protein variants. Nature Methods 4, 919–921 (2007).

17. Jumper, J. et al. Highly accurate protein structure prediction with AlphaFold. Nature 596, 583–589 (2021).

18. Lin, Z.M. et al. Evolutionary-scale prediction of atomic-level protein structure with a language model. Science 379, 1123–1130 (2023).

19. Baek, M. et al. Accurate prediction of protein structures and interactions using a three-track neural network. Science 373, 871-+ (2021).

20. Yu, T. et al. Enzyme function prediction using contrastive learning. Science 379, 1358–1363 (2023).

21. Li, F.R. et al. Deep learning-based k(cat) prediction enables improved enzyme-constrained model reconstruction. Nat Catal 5, 662-+ (2022).

22. Repecka, D. et al. Expanding functional protein sequence spaces using generative adversarial networks. Nature Machine Intelligence 3, 324–333 (2021).

23. Madani, A. et al. Large language models generate functional protein sequences across diverse families. Nat Biotechnol (2023).

24. Pucci, F., Schwersensky, M. & Rooman, M. Artificial intelligence challenges for predicting the impact of mutations on protein stability. Current opinion in structural biology 72, 161–168 (2022).

25. Liu, Y. et al. Rotamer-free protein sequence design based on deep learning and selfconsistency. Nature Computational Science 2, 451–462 (2022).

26. Dauparas, J. et al. Robust deep learning–based protein sequence design using ProteinMPNN. Science 378, 49–56 (2022).

27. Alley, E.C., Khimulya, G., Biswas, S., AlQuraishi, M. & Church, G.M. Unified rational protein engineering with sequence-based deep representation learning. Nature methods 16, 1315–1322 (2019).

28. Rives, A. et al. Biological structure and function emerge from scaling unsupervised learning to 250 million protein sequences. Proceedings of the National Academy of Sciences 118 (2021).

29. Biswas, S., Khimulya, G., Alley, E.C., Esvelt, K.M. & Church, G.M. Low-N protein engineering with data-efficient deep learning. Nature methods 18, 389–396 (2021).

30. Ofer, D., Brandes, N. & Linial, M. The language of proteins: NLP, machine learning & protein sequences. Computational and Structural Biotechnology Journal 19, 1750–1758 (2021).

31. Shin, J.E. et al. Protein design and variant prediction using autoregressive generative models. Nat Commun 12, 2403 (2021).

32. Hara, M.R. et al. S-nitrosylated GAPDH initiates apoptotic cell death by nuclear translocation following Siah1 binding. Nature cell biology 7, 665–674 (2005).

33. Tristan, C., Shahani, N., Sedlak, T.W. & Sawa, A. The diverse functions of GAPDH: views from different subcellular compartments. Cellular signalling 23, 317–323 (2011).

34. Sato, Y., Okano, K., Kimura, H. & Honda, K. TEMPURA: database of growth TEMPeratures of Usual and RAre Prokaryotes. Microbes and environments 35, ME20074 (2020).

35. Patra, S. Extremophile Protein Database. http://www.exprotdb.com/ (2018).

36. Reimer, L.C. et al. Bac Dive in 2022: the knowledge base for standardized bacterial and archaeal data. Nucleic Acids Research 50, D741–D746 (2022).

37. Costantini, S., Colonna, G. & Facchiano, A.M. ESBRI: a web server for evaluating salt bridges in proteins. Bioinformation 3, 137 (2008).

38. Perl, D., Mueller, U., Heinemann, U. & Schmid, F.X. Two exposed amino acid residues confer thermostability on a cold shock protein. Nat Struct Biol 7, 380–383 (2000).

39. Pinney, M.M. et al. Parallel molecular mechanisms for enzyme temperature adaptation. Science 371 (2021).

40. Heydenreich, F.M., Vuckovic, Z., Matkovic, M. & Veprintsev, D.B. Stabilization of G protein-coupled receptors by point mutations. Frontiers in pharmacology 6, 82 (2015).

41. Arnold, F.H. Directed Evolution: Bringing New Chemistry to Life. Angew Chem Int Ed Engl 57, 4143–4148 (2018).

42. Giver, L., Gershenson, A., Freskgard, P.O. & Arnold, F.H. Directed evolution of a thermostable esterase. Proc Natl Acad Sci U S A 95, 12809–12813 (1998).

43. Gupta, A. & Zou, J. Feedback GAN for DNA optimizes protein functions. Nature Machine Intelligence 1, 105–111 (2019).

44. Strokach, A. & Kim, P.M. Deep generative modeling for protein design. Current opinion in structural biology 72, 226–236 (2022).

45. Croitoru, F.-A., Hondru, V., Ionescu, R.T. & Shah, M. Diffusion models in vision: A survey. IEEE Transactions on Pattern Analysis and Machine Intelligence (2023).

46. Clifton, B.E., Kozome, D. & Laurino, P. Efficient exploration of sequence space by sequence-guided protein engineering and design. Biochemistry 62, 210–220 (2022).

47. Huang, P.S., Boyken, S.E. & Baker, D. The coming of age of de novo protein design. Nature 537, 320–327 (2016).

48. Wittmann, B.J., Johnston, K.E., Wu, Z. & Arnold, F.H. Advances in machine learning for directed evolution. Current opinion in structural biology 69, 11–18 (2021).

49. Dryden, D.T., Thomson, A.R. & White, J.H. How much of protein sequence space has been explored by life on Earth? Journal of The Royal Society Interface 5, 953–956 (2008).

50. Ferdinand, W. The isolation and specific activity of rabbit-muscle glyceraldehyde phosphate dehydrogenase. Biochemical Journal 92, 578 (1964).

