## supplemental information for "Protein Engineering for Thermostability through Deep Evolution"

**Supplementary Methods**

**Assessment of the** **Variant-generator**

To assess the training performance of the Variant-generator, the generator loss and discriminator loss were tracked during the training procedure. We also evaluated the similarity of generated sequences and natural sequences during the training process. The generated sequences in every 1200 steps were aligned to natural sequences to find their nearest natural sequence individually using the blastp program. After training, we further evaluated the model by analyzing the Shannon entropy of the generated sequences. The multiple sequence aliments (MSA) of 10000 generated sequences and a natural sequence of a crystallized G3PDH (PDBID: 3KV3) were constructed, and the Shannon entropy was calculated as:

$$S=-\sum_{i=1}^{20} p\left( x_{i} \right)\log_{20} p\left( x_{i} \right)$$

where p(x_i_) is the frequency of amino acids i occurring at the corresponding column based on the sequence index of the natural sequence in the MSA.

Benefiting from the GAN structure, we can use the discriminator of the GAN to evaluate the differences between the generated and natural sequences. We made box plots of the discriminant scores of the generated and natural sequences to estimate the distributional differences between them. The box plots have shown the minimum, first quartile, median, third quartile, and maximum discriminant scores of corresponding data sets.

**Assessment of the Thermo-selector model**

The losses of the Thermo-selector model in the training process were tracked to assess the training performance and to determine the end of the training procedure. After the training losses were stable, the accuracy, precision, recall, and F1 score were calculated on training and testing sets. The accuracy was calculated as:

$$Accuracy=\frac{T_{HTTP}+T_{LTTP}}{T_{HTTP}+T_{LTTP}+F_{HTTP}+F_{LTTP}}$$

where T_HTTP_ is the number of HTTP predicted as HTTP, T_LTTP_ is the number of LTTP predicted as LTTP, F_HTTP_ is the number of LTTP predicted as HTTP and F_LTTP_ is the number of HTTP predicted as LTTP.

The precision was calculated as:

$$Precision=\frac{T_{HTTP}}{T_{HTTP}+F_{HTTP}}$$

The recall was calculated as:

$$Recall=\frac{T_{HTTP}}{T_{HTTP}+F_{LTTP}}$$

And the F1 score was calculated as 2 * (Precision * Recall) / (Precision + Recall). In the building of Thermo-selector model, we have tested different parameters including the number of neural network layers and neurons. The model with best F1 score was kept for further use.

**Molecular dynamics (MD) simulations of generated and natural proteins**

The structural models of G8 and N8 were built with the AlphaFold2. The crystal structure of G3PDH form E. coli (PDBID: 1dc4) was superposed to the G8 and N8 structural models to define the binding pockets of G3P. After that the ginna^1^ docking program was used to dock the substrate G3P and coenzyme NAD to the G8 and N8 models. We build 4 scenarios, G8 at 300K, N8 at 300K, G8 at 330K and N8 at 330K to simulate. The preparation and conduction of the MD simulations were performed with the OpenMM^2^ module of python3. The simulation parameters of G3P and NAD were generated by RDkit^3^. Amber^4^ FF14 force field and tip3p water model were used to conduct all the simulations. After the default energy minimization and equilibration procedures, a 100ns productive simulation was run for each scenario. The prody^5^ and pylab^6^ modules were used to analyze the simulation trajectories. The root mean square fluctuation (RMSF) of last 50ns of each simulation were calculated to compare the overall flexibility of G8 and N8 at different temperatures. This metric indicates the average deviation of a particle over time from a reference position.

**MDH activity and thermal stability assay**

To test MDH activity and thermal stability, proteins with 0.1 M potassium phosphate buffer (pH 7.4) were held at 30°C or 65°C for 15 min, and then added to a reaction mixture containing 0.15 mM NADH, 0.2 mM oxaloacetic acid and 0.1 M potassium phosphate buffer (pH 7.4). The final reaction volume was 100 μL and the reaction was carried out at room temperature in a UV-transparent 96-well half-area plate (UV-Star Microplate, Greiner). Activity was measured in triplicates by following NADH oxidation to NAD^+^, with an absorbance reading at 340 nm performed every 30s for 30 min in a BMG Labtech SPECTROstar Nano spectrophotometer. Unspecific oxidation of NADH was monitored in no-substrate controls, and these values were subtracted from the other samples. Conversion from absorption values to NADH concentration was carried out using an extinction coefficient of 6.22 mM. The activity of MDH were calculated with those formula Units/mg/min = △A340 x VT(Volume of tube)/(6.22 x Concentration(mg) x Time(s)).

**Determination of protein concentration**

Protein concentrations were determined using Pierce™ BCA protein detection kit (Thermo Fisher, 23227). According to the detection degree of SDS-PAGE, the protein was diluted by a certain number of times. 25μL protein solution was taken into the 96-well plate, then 200μL mixture of liquid A and liquid B (50:1) was added. The plate was placed at 37℃ for 30min, and the absorbance at λ=562 nm was detected by enzymoleter, and the OD value under this absorbance was recorded. The recorded OD value is substituted into the formula of the standard curve drawn, and the result is multiplied by the corresponding dilution multiple to obtain the concentration of the purified protein sample.

**Assembly and correction of generated gene sequences**

50~60nt short ssDNA are synthesized by Zhong He Gene Co.Ltd(Tianjin). 20-30 ssDNA fragments, primers, Pfu mixture were mixed to make a PCA reaction system. After several PCR cycles, gene sequences were assembled. After that, mismatch repair enzyme(commercial) was added to the system and the reaction was carried out at 37℃ for 2 hours to correct the assembly error. Assembled and corrected genes were cloned into expected plasmid and transfected into competent cells.

**Supplementary Table**

Supplementary Table 1 | Overview and assessment of experimental results. Protein solubility was assessed based on SDS-page gels (sourcedata) and activity was assessed based on measurements of G3PDH activity.

| **Protein ID** | **Solubility** | **Competent Cell** | **Activity (30℃)** | **Activity (65℃)** |
| --- | --- | --- | --- | --- |
| G3PDH | S | BL21(DE3) | 34.8±2.62 | - |
| TDH1 | S | BL21(DE3) | 32.6±1.87 | - |
| G1 | S | BL21(DE3) | 9.4±0.10 | - |
| G2 | S | BL21(DE3) | 53.6±0.13 | - |
| G3 | S | BL21(DE3) | 59.6±0.57 | - |
| G4 | U | - | - | - |
| G5 | U | - | - | - |
| G6 | U | - | - | - |
| G7 | S | BL21(DE3) | 51.9±2.68 | 44.4±1.95 |
| G8 | S | Arctic Express (DE3) | 46.5±3.90 | 38.9±0.67 |
| G9 | S | Arctic Express (DE3) | 34.8±2.45 | 2.6±0.08 |
| G10 | S | Arctic Express (DE3) | 32.7±0.14 | 23.9±0.48 |
| G11 | S | BL21(DE3) | 31.5±0.66 | 41.3±0.26 |
| G12 | S | BL21(DE3) | 29.9±1.27 | 11.7±0.12 |
| G13 | S | BL21(DE3) | 19.7±0.20 | 6.6±0.80 |
| G14 | S | BL21(DE3) | 19.2±1.73 | 15.1±1.31 |
| G15 | S | BL21(DE3) | 14.0±1.02 | 22.4±1.32 |
| G16 | S | Arctic Express (DE3) | 9.5±0.01 | 1.5±0.15 |
| G17 | S | Arctic Express (DE3) | 9.4±0.17 | 3.8±0.06 |
| G18 | S | Arctic Express (DE3) | 5.6±0.15 | - |
| G19 | S | Arctic Express (DE3) | 5.4±0.01 | - |
| G20 | S | BL21(DE3) | 5.2±0.15 | - |
| G21 | S | Arctic Express (DE3) | 4.3±0.03 | - |
| G22 | S | Arctic Express (DE3) | 2.7±0.07 | - |
| G23 | S | Arctic Express (DE3) | 2.4±0.02 | - |
| G24 | S | BL21(DE3) | - | - |
| G25 | S | Arctic Express (DE3) | - | - |
| G26 | S | BL21(DE3) | - | - |
| G27 | S | BL21(DE3) | - | - |
| G28 | S | Arctic Express (DE3) | - | - |
| G29 | S | BL21(DE3) | - | - |
| G30 | U | - | - | - |
| G31 | U | - | - | - |
| G32 | U | - | - | - |
| G33 | U | - | - | - |
| G34 | U | - | - | - |
| G35 | U | - | - | - |
| G36 | U | - | - | - |
| N1 | S | BL21(DE3) | 9.1±0.53 | - |
| N2 | S | BL21(DE3) | 62.2±0.08 | 14.1±1.34 |
| N3 | S | BL21(DE3) | 60.3±1.33 | 14.9±0.77 |
| N7 | S | BL21(DE3) | 26.7±2.37 | - |
| N8 | S | BL21(DE3) | 2.0±0.22 | - |
| N10 | S | BL21(DE3) | 29.8±0.73 | - |
| N11 | S | BL21(DE3) | 31.9±1.40 | 2.5±0.02 |
| N12 | S | BL21(DE3) | 24.1±3.10 | 2.1±0.04 |
| N13 | S | BL21(DE3) | 17.1±0.67 | 10.7±1.40 |

*Asterisk represents the level of enzyme activity

Supplementary Table 2 Experimental tested HTTP compared with their nearest natural sequence.

| Generated | Natural | Identity | Substitutions | NCBI ID | Natural sequence OTG |
| --- | --- | --- | --- | --- | --- |
| G1 | N1 | 93.114 | 23 | WP_100548227.1 |  |
| G2 | N2 | 92.836 | 24 | WP_108039253.1 |  |
| G3 | N3 | 87.915 | 40 | WP_036936343.1 |  |
| G4 | N4 | 74.107 | 87 | OGI98331.1 |  |
| G5 | N5 | 64.923 | 111 | WP_014797236.1 |  |
| G6 | N6 | 59.701 | 131 | WP_021660543.1 |  |
| G7 | N7 | 92.192 | 26 | WP_093976896.1 | Low |
| G8 | N8 | 90.332 | 31 | WP_018610755.1 | Low |
| G9 | N9 | 94.26 | 19 | WP_120258102.1 |  |
| G10 | N10 | 87.387 | 42 | WP_045802628.1 | Median |
| G11 | N11 | 91.892 | 27 | WP_126020941.1 | Median |
| G12 | N12 | 85.241 | 49 | WP_072755848.1 | high |
| G13 | N13 | 93.393 | 22 | WP_027077543.1 | Low |
| G14 | N7 | 93.694 | 21 | WP_093976896.1 | Low |
| G15 | N7 | 93.694 | 21 | WP_093976896.1 | Low |
| G16 | N16 | 86.787 | 44 | WP_106147095.1 |  |
| G17 | N17 | 88.485 | 38 | WP_040249034.1 |  |
| G18 | N18 | 76.506 | 77 | WP_014759316.1 |  |
| G19 | N18 | 78.443 | 72 | WP_014759316.1 |  |
| G20 | N20 | 88.889 | 37 | WP_112239825.1 |  |
| G21 | N10 | 85.886 | 47 | WP_045802628.1 |  |
| G22 | N22 | 87.087 | 42 | WP_161649912.1 |  |
| G23 | N23 | 72.121 | 92 | WP_109826737.1 |  |
| G24 | N24 | 83.533 | 55 | WP_015256252.1 |  |
| G25 | N25 | 84.894 | 49 | WP_071111430.1 |  |
| G26 | N24 | 85.329 | 49 | WP_015256252.1 |  |
| G27 | N24 | 86.826 | 43 | WP_015256252.1 |  |
| G28 | N24 | 86.826 | 43 | WP_015256252.1 |  |
| G29 | N8 | 86.486 | 45 | WP_018610755.1 |  |
| G30 | N30 | 91.291 | 29 | WP_179242944.1 |  |
| G31 | N31 | 77.477 | 73 | WP_013787603.1 |  |
| G32 | N32 | 90.691 | 31 | WP_047245810.1 |  |
| G33 | N24 | 87.126 | 43 | WP_015256252.1 |  |
| G34 | N20 | 83.384 | 55 | WP_112239825.1 |  |
| G35 | N11 | 94.595 | 18 | WP_126020941.1 |  |
| G36 | N36 | 77.946 | 73 | WP_132769025.1 |  |

Supplementary Table 3 Mutations in verified HTTPs compared to their individual nearest natural sequence. Single change which no contact with other mutation and changes in cluster are list separately. Members in the same cluster are connected with ‘+’ sign.

| **Generated sequence ID** | **single change to natural sequence** | **changes in cluster to natural sequence** |
| --- | --- | --- |
| G7 | 185LEU, 255ASP, 294VAL, 305ASN | 3SER+26GLY+29GLU+24LYS, 47LYS+54ASN+56ASP, 125LYS+126ASP, 166ASN+260VAL, 214VAL+217GLY+222ALA+224LYS, 286VAL+317ALA+289SER, 329HIS+330VAL+331ALA+332SER |
| G8 | 6ILE, 63ASP, 79ILE, 88GLY, 113LYS, 140SER, 181THR, 232SER, 262MET, 270LEU, 283ALA, 331VAL | 2SER+3ILE, 23GLN+25PHE+27ASN, 48LYS+60ILE+69VAL+57GLN+59ASP, 164LEU+167LYS+168TRP+250VAL, 256ASP+257GLU, 278GLU+298ASP, 289ASP+291ARG+294ILE+312ALA |
| G10 | 3LYS, 54ARG, 79LYS, 83HIS, 105ASP, 142THR, 146VAL, 170VAL, 195TRP, 231SER, 237VAL, 294VAL, 305ASN | 6LEU+93TYR, 21ALA+317TYR+287SER+289SER, 65ASN+67PHE+71HIS+73ILE+75VAL, 137LYS+138GLU, 164HIS+166ALA+260VAL+263LYS+267THR+268GLU+255ASP+259ALA, 202LEU+203LEU, 217GLY+222SER+224LYS, 328ILE+330ILE+332GLY |
| G11 | 41GLU, 48ARG, 65ASN, 78ALA, 89ASP, 107LYS, 155VAL, 226GLY, 256ALA, 283SER | 5VAL+96VAL+121MET, 21ILE+22ILE, 59THR+61GLU+68VAL, 83SER+84ASN, 261ALA+265ALA+269PRO, 278ASP+279GLU+298GLY+300ALA |
| G12 | 23GLU, 39ASP, 80LYS, 107LEU, 193LYS, 219LYS, 226GLY, 256GLU, 283SER, 312SER | 2MET+27ASN, 5LEU+93ASP+96VAL+120ILE+324VAL+328VAL+331ILE+333LYS, 16MET+18LEU, 48LYS+55ALA, 61SER+68ILE+73LYS+64GLY+65ASN+66PHE, 88ASN+89GLU+90VAL, 112LEU+113LYS, 138ASP+139LYS+140SER, 161SER+165ASN+166GLU, 251LYS+252GLY, 261ALA+264GLU+265ALA+269SER, 279GLU+280SER+298LYS |
| G14 | 47LYS, 56ASP, 105ASP, 166ALA, 172GLY | 3LYS+26GLY+27ASP+29GLU+24LYS, 125LYS+126ASP, 222ALA+224GLU, 286VAL+317TYR, 327ILE+329HIS+330VAL+331ALA+332GLY |
| G15 | 47LYS, 56ASP, 84ILE, 88ALA, 138ASP, 263GLU | 22CYS+24LYS+26GLY, 125LYS+126ASP, 222ALA+224LYS, 286VAL+317ALA, 323MET+327ILE+329HIS+330VAL+331ALA+332GLY |

**Supplementary Figures**

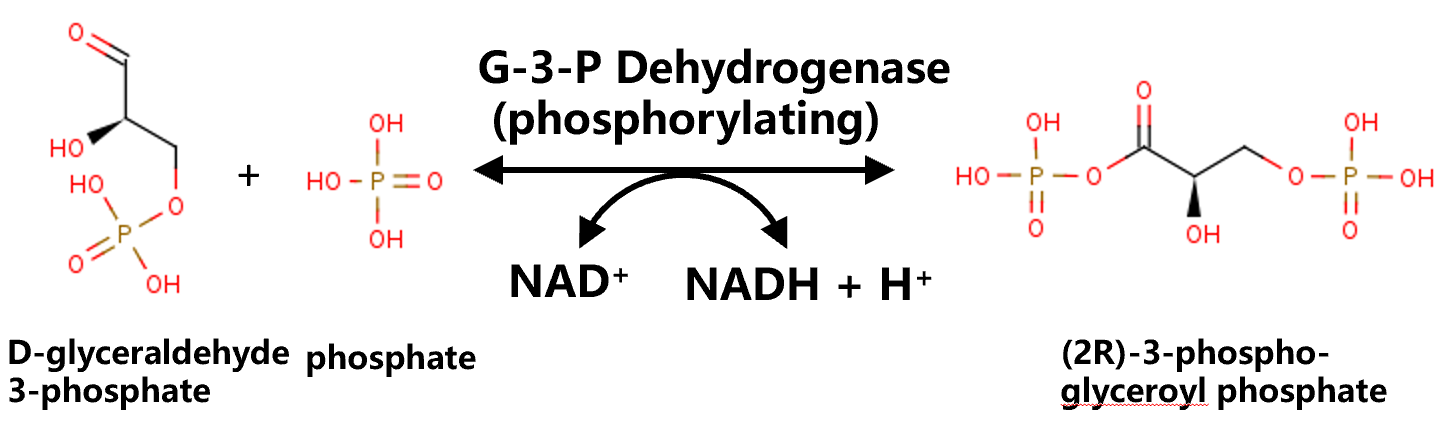

**Figure S1 The reaction catalyzed by G3P Dehydrogenase (EC 1.2.1.12).** G3PDH activities is measured by determining the increase of NADH.

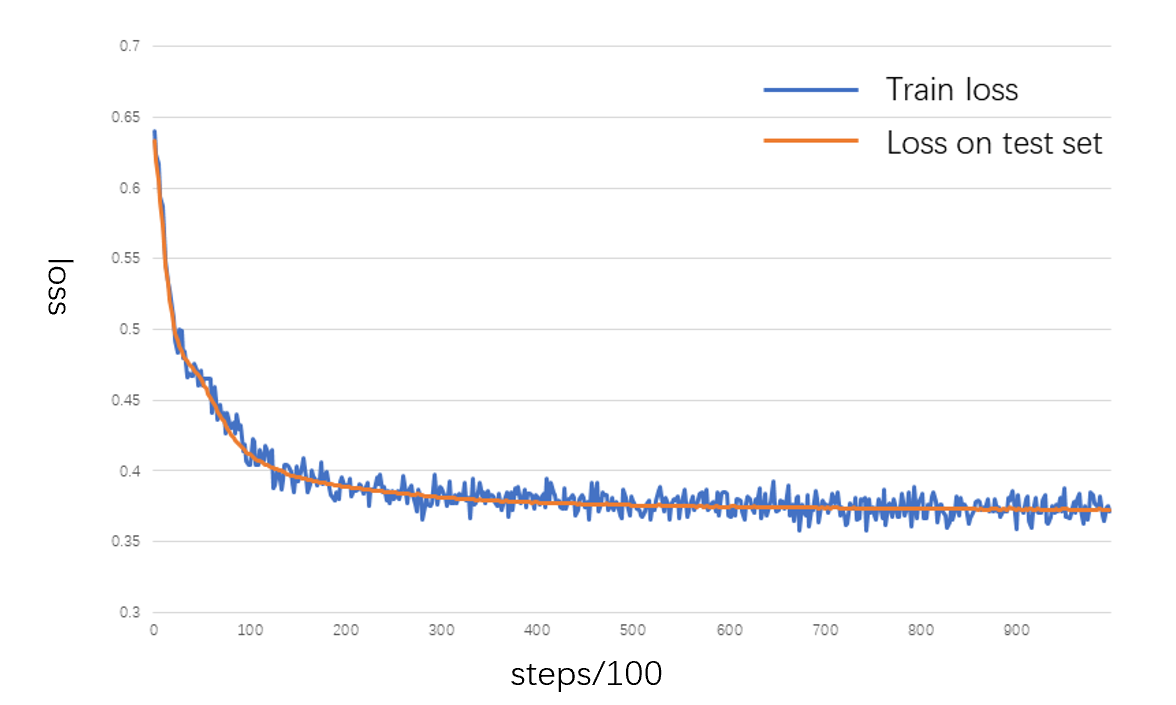

**Figure S2 The train loss and loss on test set during the training of Thermo-selector.** The train loss are calculated on a batch and the loss on test set are calculated on the whole set.

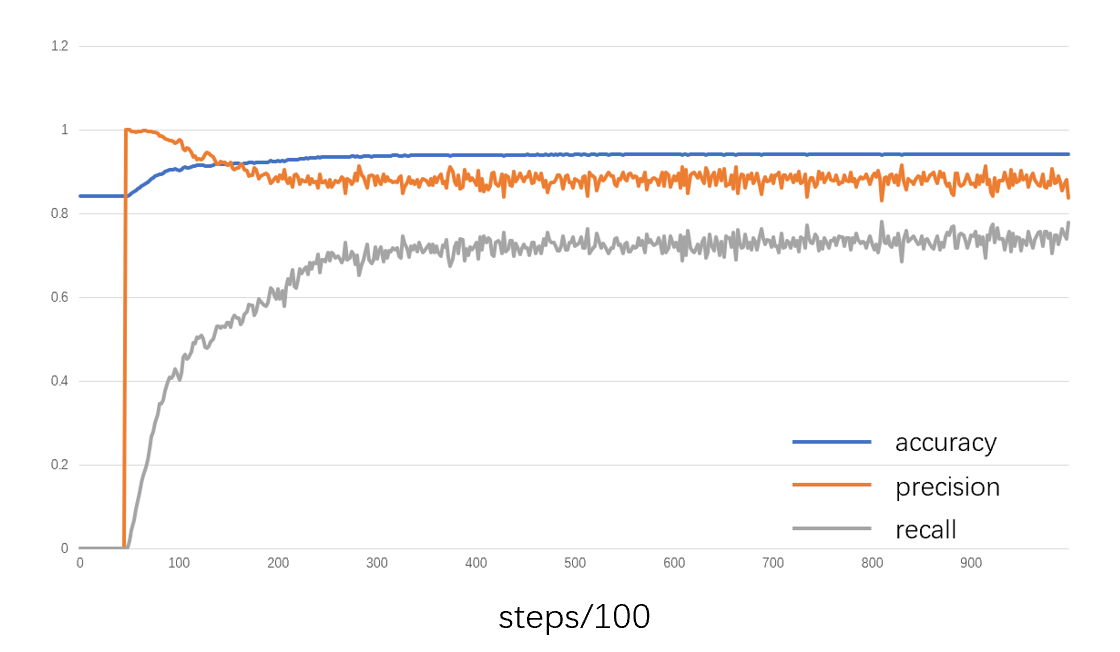

**Figure S3 Accuracy, precision, and recall of HTTPs on test set during the training of Thermo-selector.** All metrics are calculated on the whole set.

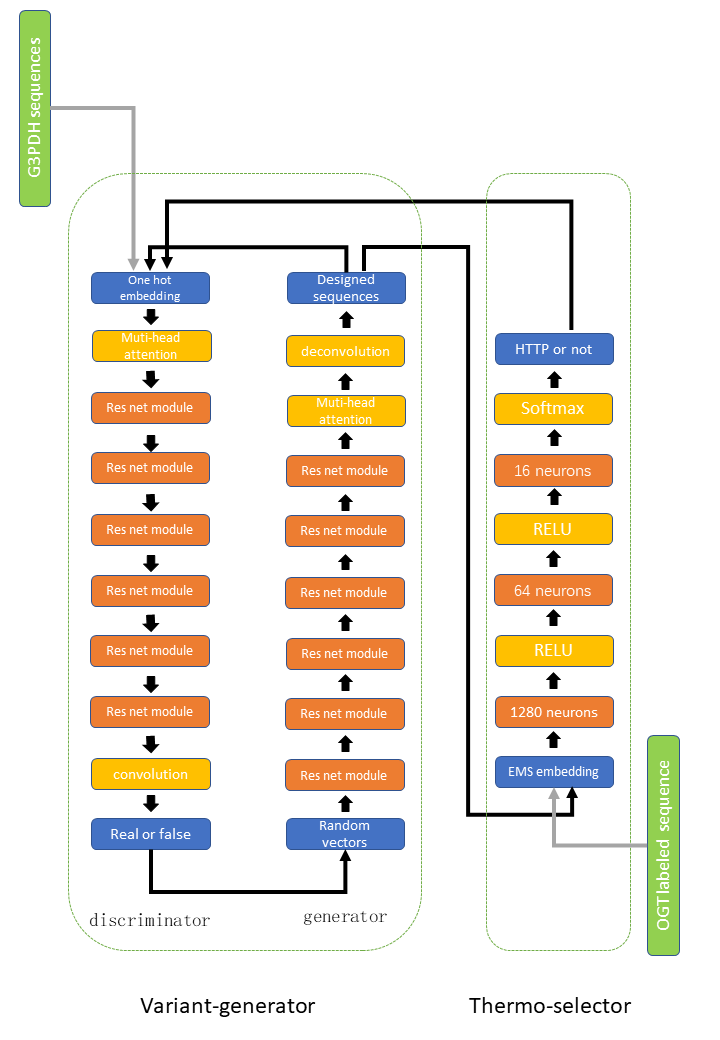

**Figure S4 Model structure of Variant-generator and Thermo-selector**. The structure of Variant-generator is based on ProteinGAN and the Thermo-selector utilized ESM as an encoder.

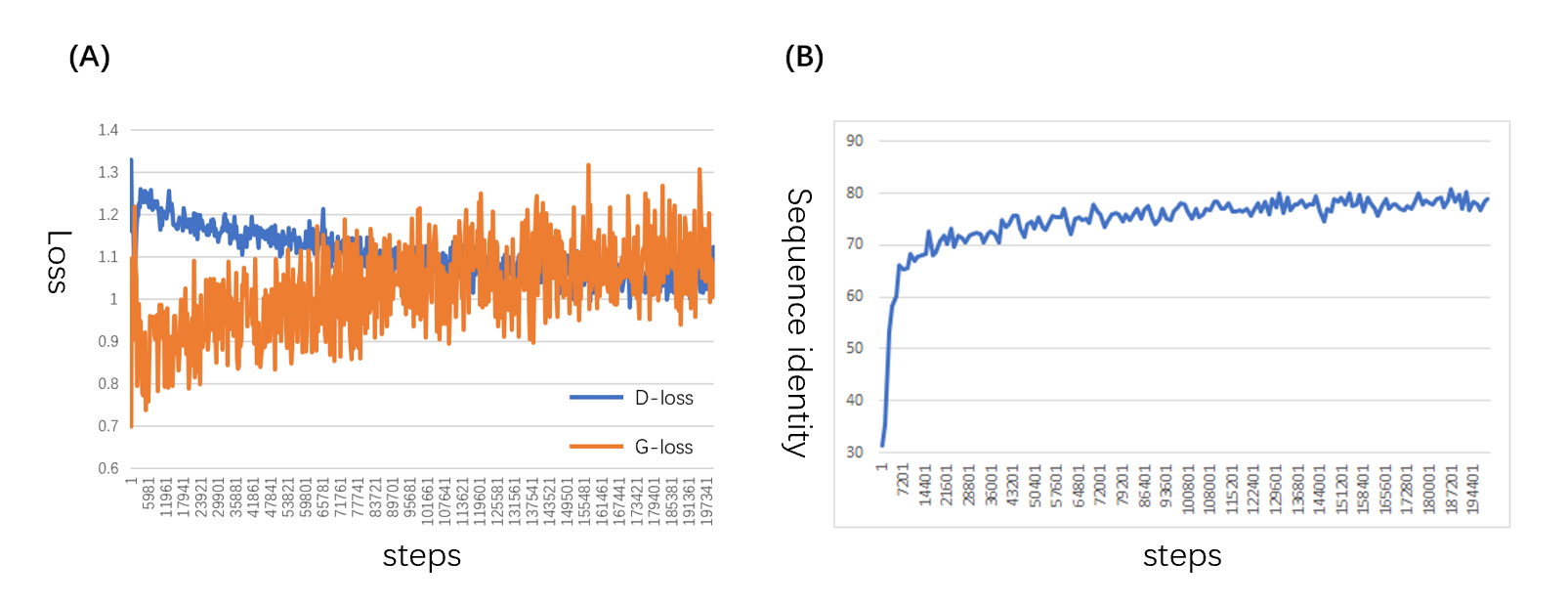

**Figure S5 Evaluating the training process of Variant-generator.** (A)generator loss(G-loss) and discriminator loss(D-loss) during the training of the GAN based Variant-generator. (B) Average generated sequence identity to natural sequences during the training of the Variant-generator.

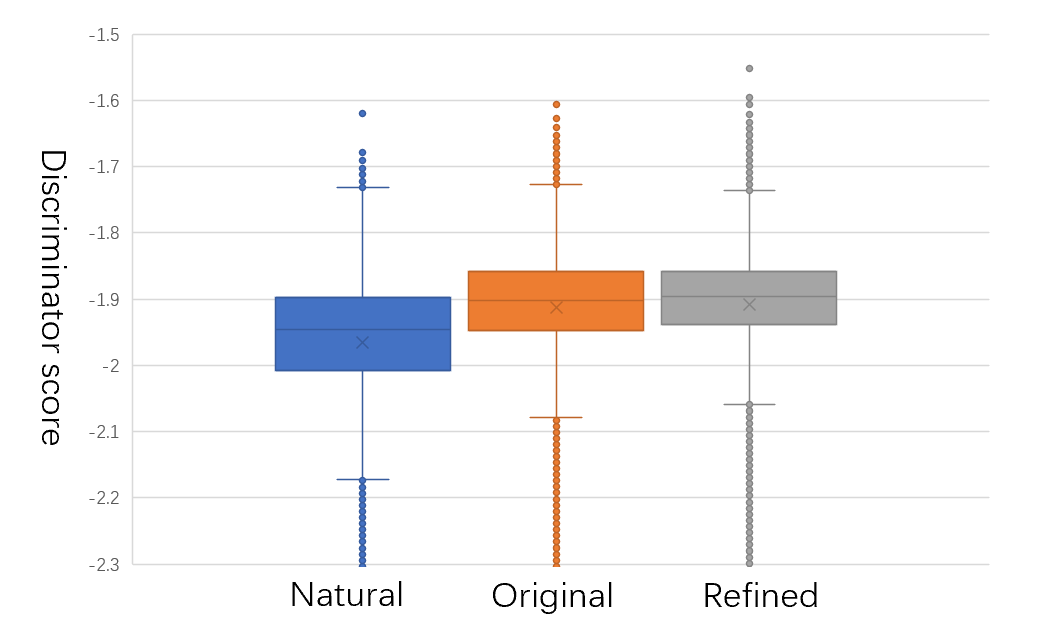

**Figure S6. Distribution of discriminator scores.** The distribution of discriminator score of natural sequences(blue), original Variant-generator generated sequences(orange), refined Variant-generator generated sequences(grey) are shown. All the scores are calculated by the discriminator of the GAN structure from the original Variant-generator.

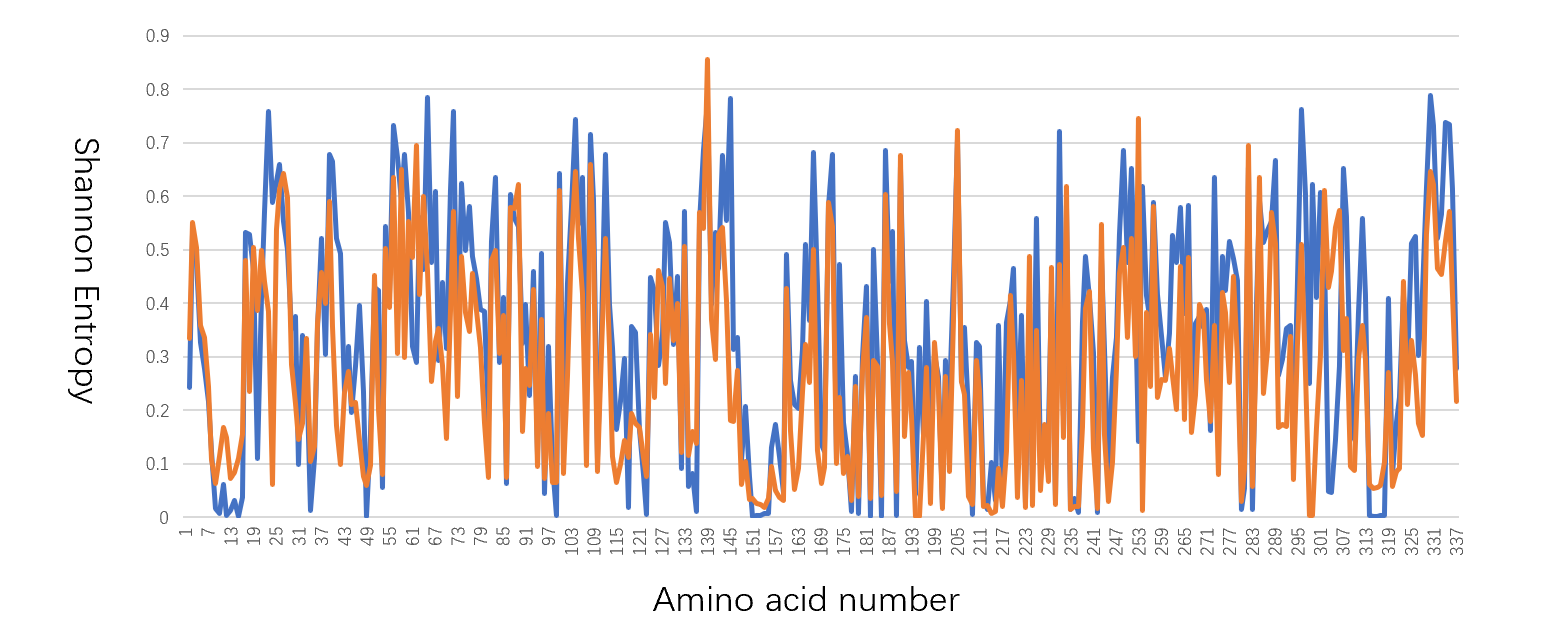

FigureS7. **The Shannon entropy of natural and generated sequences**. The Shannon entropy of natural sequences(blue) and original generated sequences(orange) are shown. All the sequences are aligned to the sequence of a crystallized G3PDH (PDBID: 3KV3).

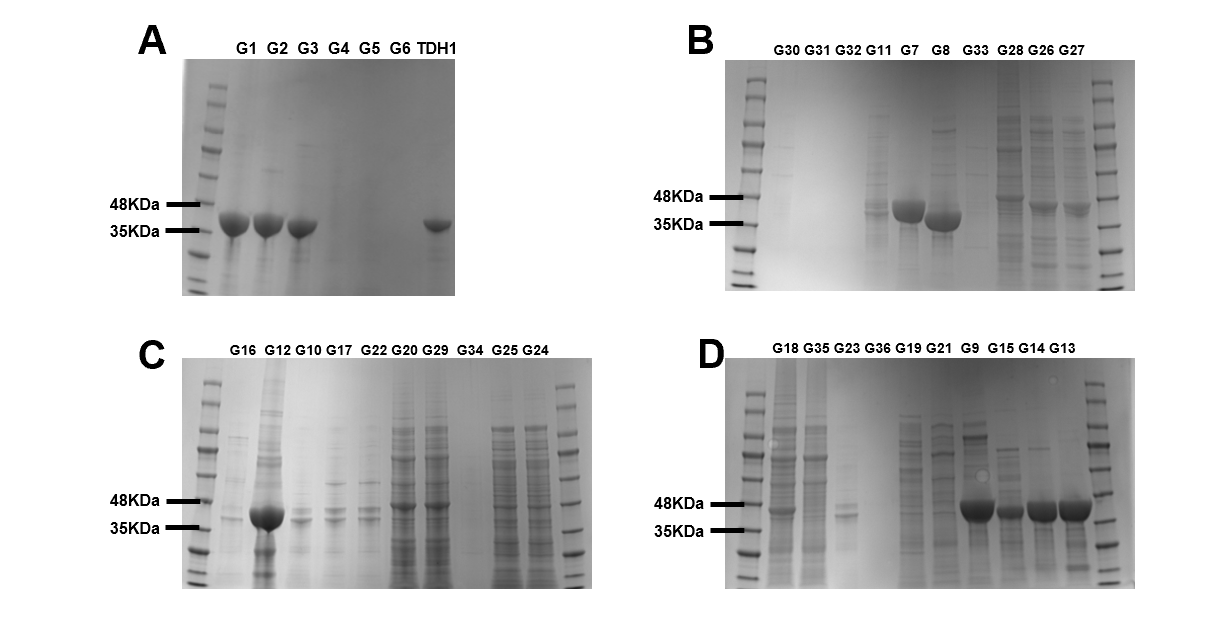

**Figure S8 SDS-PAGE of correctly folded generate sequences, commercial G3PDH as control.**

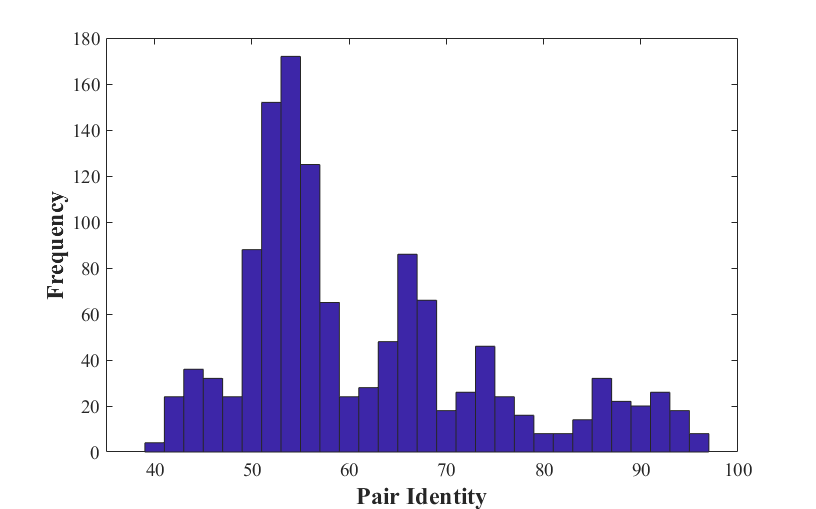

Min = 40

Max = 96

Mean = 61

Median = 56

**Figure S9. Distribution of blast pair identities among 30 generated G3PDH sequences.**

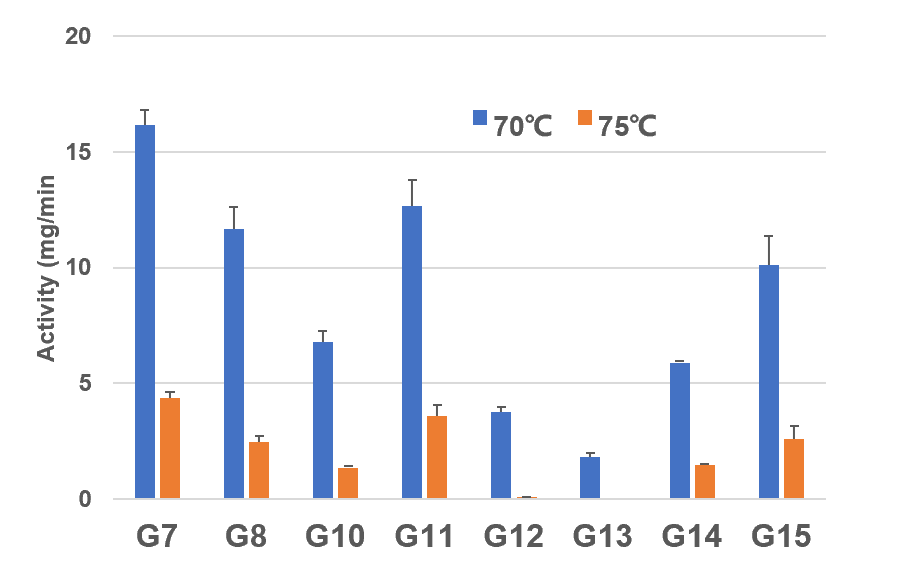

**Figure S10. Thermal stability assay of 8 generated G3PDH’s HTTPs at 70℃ and 75℃.** All 8 proteins showed enzyme activities at 70 ℃, 6 proteins (i.e., G7, G8, G10, G11, G14, G15) showed enzyme activities at 75 ℃.

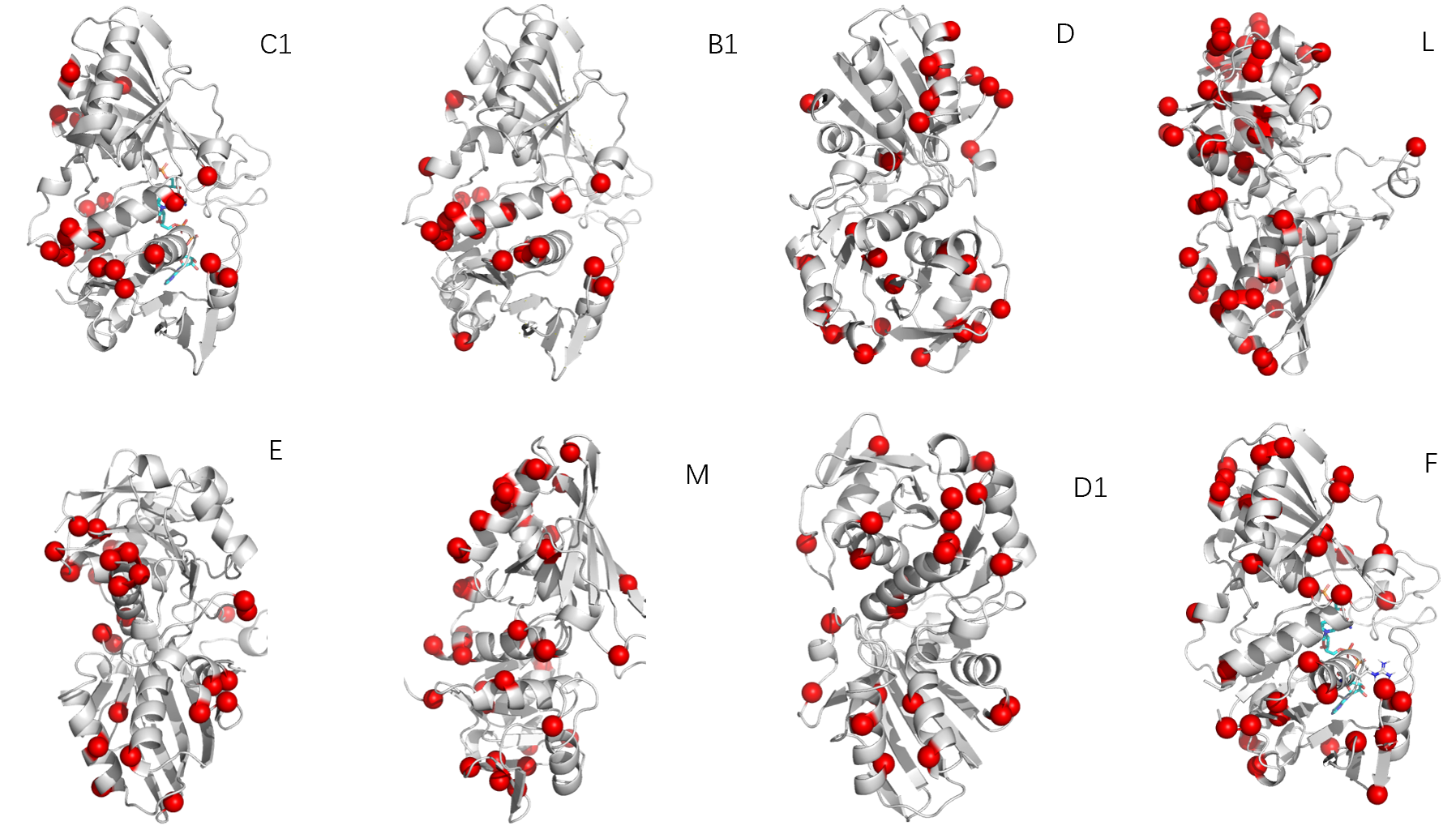

**Figure S11 Structure models of verified HTTPs.** Mutations compared to their individual nearest natural protein are shown as red dots. The structure figures were made by pymol^7^.

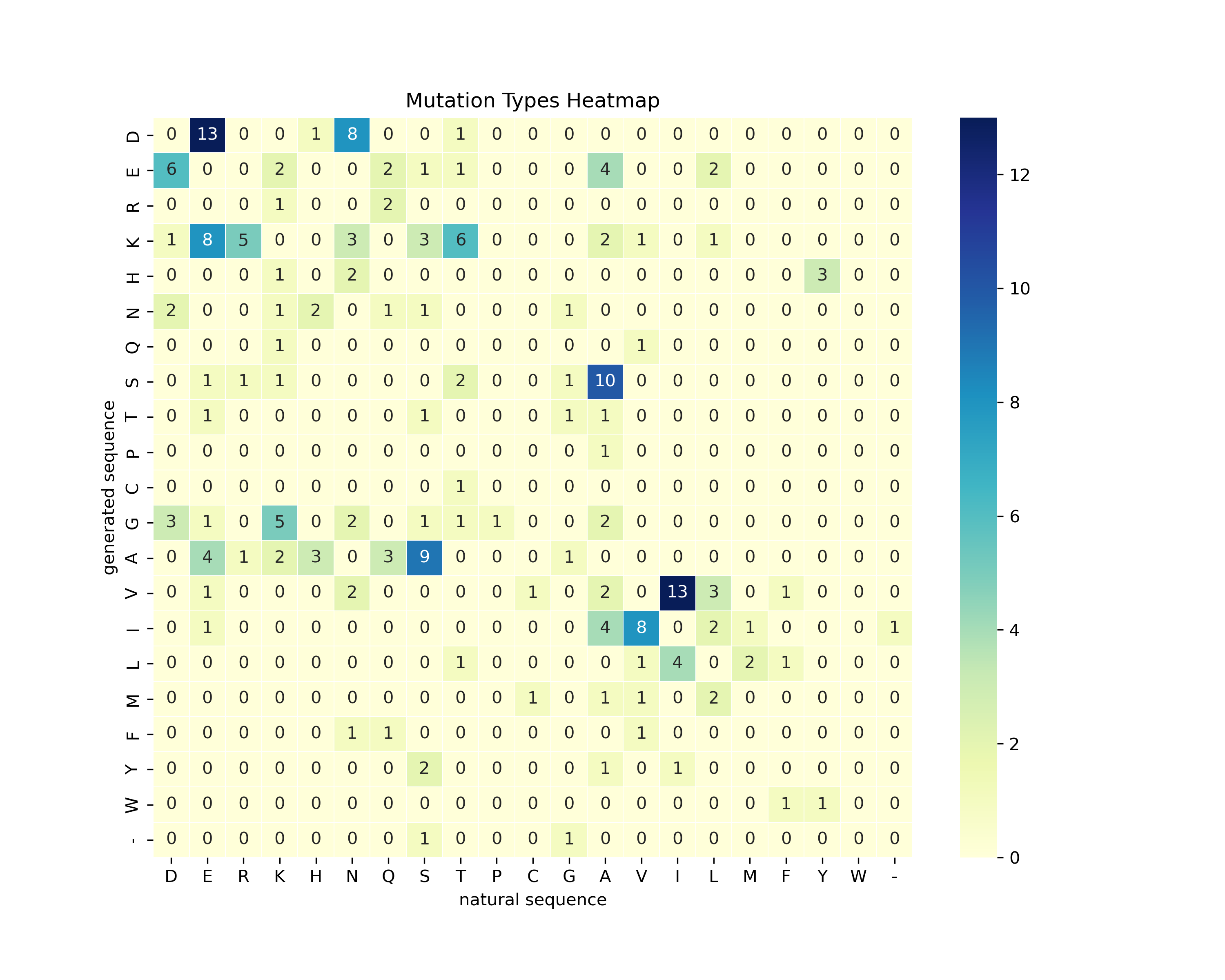

**Figure S12 The heatmap of different mutation types of verified HTTPs with their nearest natural sequences.** The numbers of each type of mutation are shown in the figure.

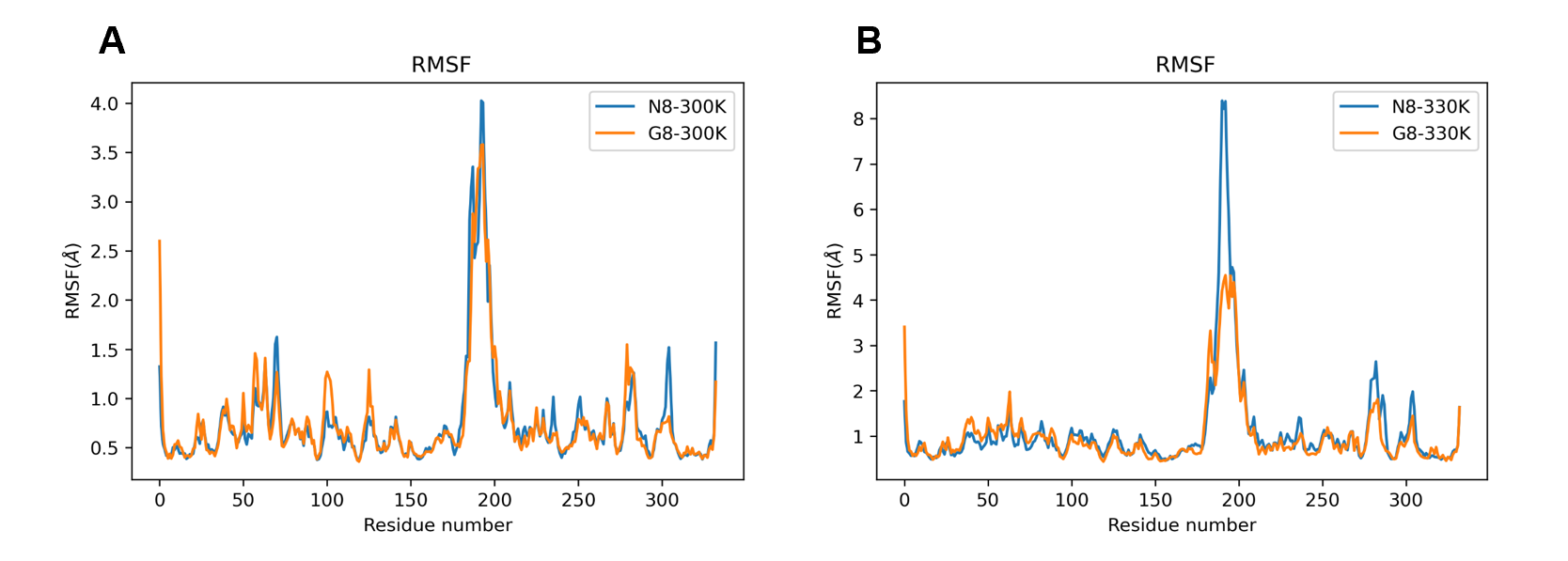

**Figure S13 Figure S13 RMSF of G8 and N8 from MD simulations at different temperature.** (A) RMSF of G8(orange) and N8(blue) at 300K. (B) RMSF of G8(orange) and N8(blue) at 330K.

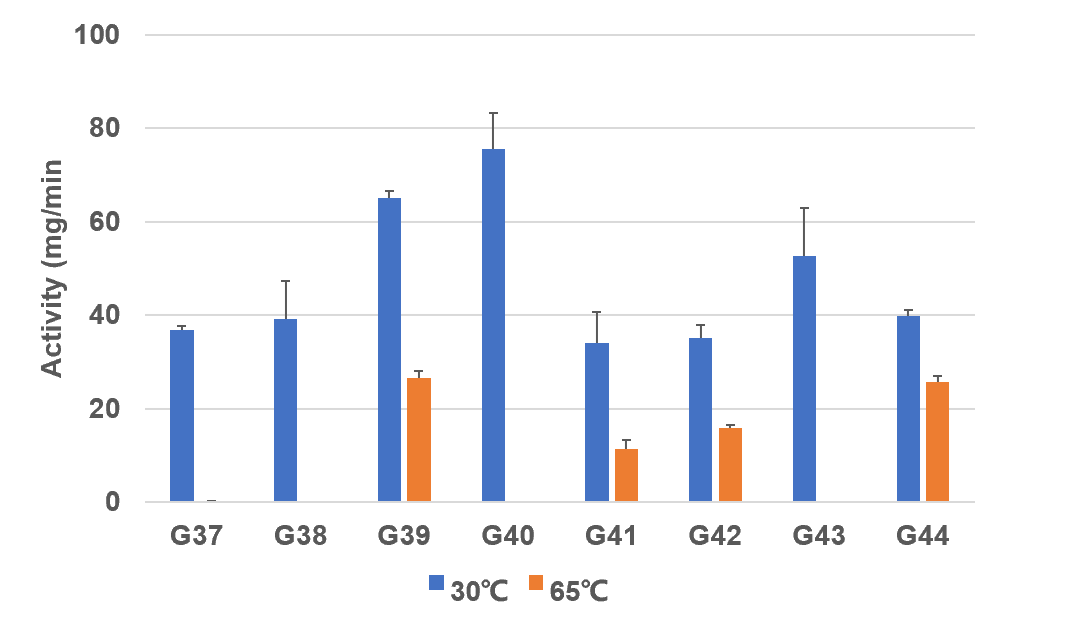

**Figure S14 G3PDH assay of proteins that only filtered by Thermo-selector, commercial G3PDH as control.**

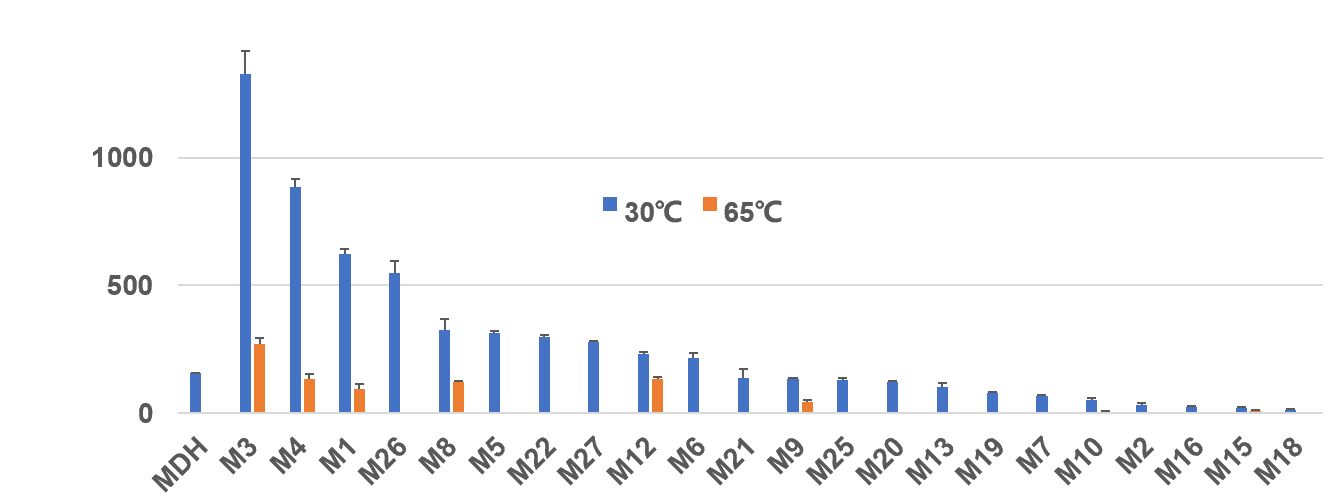

**Figure S15 MDH enzyme activity assay of 22 generated and expressed proteins.** Commercial MDH as control. All 22 protein showed enzyme activities at 30 ℃, 6 proteins (i.e., M3, M4, M1, M8, M12, M9) showed enzyme activities at 65 ℃.

**Supplementary references**

1.	McNutt, A.T. et al. GNINA 1.0: molecular docking with deep learning. *Journal of cheminformatics* **13**, 1-20 (2021).

2.	Eastman, P. et al. OpenMM 7: Rapid development of high performance algorithms for molecular dynamics. *PLoS computational biology* **13**, e1005659 (2017).

3.	Landrum, G. RDKit: A software suite for cheminformatics, computational chemistry, and predictive modeling. *Greg Landrum* **8** (2013).

4.	Case, D.A. et al. Amber 2021. (University of California, San Francisco, 2021).

5.	Bakan, A., Meireles, L.M. & Bahar, I. ProDy: protein dynamics inferred from theory and experiments. *Bioinformatics* **27**, 1575-1577 (2011).

6.	Hunter, J.D. Matplotlib: A 2D graphics environment. *Computing in science & engineering* **9**, 90-95 (2007).

7.	DeLano, W.L. Pymol: An open-source molecular graphics tool. *CCP4 Newsl. Protein Crystallogr* **40**, 82-92 (2002).
